# Layer-specific input to medial prefrontal cortex is linked to stress susceptibility

**DOI:** 10.1101/2024.04.03.587746

**Authors:** Sabahaddin Taha Solakoğlu, Şefik Evren Erdener, Olga Gliko, Alp Can, Uygar Sümbül, Emine Eren-Koçak

## Abstract

Stress response is essential for adapting to an ever-changing environment. However, the mechanisms that render some individuals susceptible to stress are poorly understood. While chronic stress is known to induce dendritic atrophy and spine loss in medial prefrontal cortex (mPFC), its impact on synapses made by long-range projections terminating on the mPFC remains unknown. Here, we labeled synapses on male mouse mPFC dendrites formed by ventral hippocampus (VH), basolateral amygdala (BLA) and ventral tegmental area (VTA) long-range afferents using different-colored eGRASP constructs. We obtained multispectral 3D-images of the mPFC covering all cortical laminae, and automatically segmented the dendrites and synapses. In layer II/III, the relative abundances and spatial organizations of VH-mPFC and BLA-mPFC synapses changed similarly in stress resilient (SR) and stress susceptible (SS) mice when compared to stress naïve (SN) mice. In layers Vb and VI, on the other hand, the percentage of BLA-mPFC increased and that of VH-mPFC decreased only in SS mice. Moreover, the distances of VH synapses to their corresponding closest BLA synapses decreased and the distances of BLA synapses to their corresponding closest VH synapses increased in the SS group. Consistently, the percentage of single dendritic segments receiving input from multiple brain regions increased in the SS group, suggesting that long-range synaptic inputs to deep layers of mPFC were disorganized in SS mice. Our findings demonstrate afferent- and lamina-specific differential reorganization of synapses between different stress phenotypes, suggesting specific roles for different long-range projections in mediating the stress response.

## INTRODUCTION

Stress is an essential adaptive response to imminent or perceived changes in the environment to restore homeostasis. In some individuals, however, stress response can go awry resulting in mental disorders like depression, anxiety disorders and posttraumatic stress disorders for yet unknown reasons. Understanding mechanisms underlying stress susceptibility, therefore, is important for developing prevention and early intervention strategies against stress-related mental disorders.

A key region mediating the stress response is the medial prefrontal cortex (mPFC). Chronic stress causes atrophy of apical dendrites and loss of spines in layer II/III and layer V of mPFC pyramidal neurons without changes in basal dendrites (1–6). These effects of chronic stress are circuit specific: entorhinal cortex-projecting pyramidal neurons show dendritic atrophy in response to chronic stress, whereas basolateral amygdala (BLA)-projecting pyramidal neurons are resilient to stress-induced dendritic remodeling (7). Whether such dendritic resilience to remodeling also applies to afferent projections to the mPFC remains to be answered. Chronic stress-induced dendritic atrophy would be expected to result in the loss of synaptic connections that these dendrites make. If long-range projections to the mPFC from various brain regions are affected differently by this synaptic loss, it will alter the relative weight of inputs from different brain regions on the mPFC and thus, alter the brain circuitry involved in stress response. To date, however, **which mPFC-afferents are affected by dendritic retraction or spine loss is unknown**. Similarly, **whether the reported morphological changes in the dendritic arbor are adaptive or maladaptive remains unknown, because susceptibility to stress was not evaluated in the previous literature**.

mPFC receives input from ventral hippocampus (VH), basolateral amygdala (BLA) and ventral tegmental area (VTA); brain regions indicated in the regulation of the stress response (8, 9). Optogenetic or chemogenetic activation of mPFC or mPFC-projecting VTA, VH and BLA neurons were reported to alter depression- and anxiety-like behaviors (10–19). Unlike sensory cortices, mPFC is agranular, lacks a thalamus projecting layer IV, and receives inputs across all layers (8). Information from various long-range afferents into different layers is integrated by the local circuits within the same layer (intralaminar) or across different layers (interlaminar) to generate an optimal response to adapt to everchanging conditions. **Little is known on how chronic stress alters mPFC circuitry across cortical laminae,** as earlier studies mostly focused on single layers (20). Evidence indicates that long-range inputs may regulate local circuits selectively since they preferentially target certain cortical layers: Layer II/III and layer V for BLA and VH, respectively (8, 21–23). Distinct mPFC projecting neuron populations identified in both BLA (fear and extinction neurons) and VH (approach and avoidance neurons) regulate stress response and related fear in opposite directions; promoting or suppressing it by integrating the perceptual, emotional and spatial data arriving at mPFC (24, 25). Whether these neurons with opposite functions project to the same or different layers of mPFC is not known. Examining chronic stress-induced changes in specific synaptic inputs across all layers of the mPFC is therefore, crucial to understanding how stress alters brain circuits to produce adaptive responses, as well as what goes awry in stress-related mental disorders.

In this study, we investigated the differences in synapses on the mPFC originating from the VH, BLA, and VTA after stress exposure in stress-resilient (SR) and stress-susceptible (SS) mice, examining all cortical layers. For this purpose, we acquired multispectral, high resolution 3D images of a ∼200 × 1500 × 30 μm^3^ volume from the mPFC and mapped VH-mPFC, BLA-mPFC, and VTA-mPFC synapses across all layers of the cortical column using the enhanced green fluorescent protein reconstitution across synaptic partners (e-GRASP) system (26). We hypothesized that the relative abundance of VH-mPFC, BLA-mPFC and VTA-mPFC synapses would change following stress exposure, but differently in stress resilient (SR) and stress susceptible (SS) mice. Specifically, we predicted that, following stress exposure, the relative abundance of BLA-mPFC and VTA-mPFC synapses would increase, while the relative abundance of VH-mPFC synapses will decrease in SS mice compared to the SR group.

## MATERIALS AND METHODS

### Animals

Wild-type male C57BL/6 mice (12–16-week-old) and wild-type CD-1 male mice (4–20-month-old) were used (Kobay AS and Hacettepe University Laboratory Animals Research and Application Centre). Mice were maintained at 12:12 light:dark cycle (07:00-19:00) and stable temperature (19-22 °C) with *ad libitum* access to food and water. All procedures were done in accordance with Animal Experimentations Local Ethics Board of Hacettepe University (Ethics number: 2019/78).

### Enhanced GFP (green fluorescent protein) Reconstitution Across Synaptic Partners (e-GRASP) system and Construction of Adeno Associated Virus

We used enhanced green fluorescent protein (GFP) reconstitution across synaptic partners (e-GRASP) system to differentially label functional synapses between VH, VTA and BLA excitatory projections and mPFC pyramidal neurons (27–29). The e-GRASP system enables the visualization of functional synapses through the complementation of presynaptic neurexin and postsynaptic neuroligin. Each protein is fused with complementary fragments of split GFP, so when neurexin binds to neuroligin, GFP is reconstituted without significantly altering their binding affinities (27–30). The plasmids pAAV-CWB-yellow-pre-eGRASP(p32), pAAV-CWB-cyan-pre-eGRASP(p30), pAAV-CWB-green-pre-eGRASP(p32), pAAV-EWB-DIO-myrTagRFP-T-P2A-post-eGRASP were gifts from Bong-Kiun Kaang (Addgene plasmids #111580, #111581, #111586, #111598) (26) and packaged into AAV 2/1 at the Viral Core Facility of Charite-Universtatmedizin Berlin. AAV2/1-CaMKIIα-iCre was obtained from the same facility.

### Intracranial Virus Injections

Mice were anesthetized with ketamine [50-100 mg/kg, intraperitoneal (ip)] and xylazine (5-10 mg/kg, ip), then maintained under 1-2% isoflurane or sevoflurane anesthesia. AAV2/1-pAAV-CWB-yellow-pre-eGRASP (p32), AAV2/1-pAAV-CWB-green-pre-eGRASP (p32) and AAV2/1-pAAV-CWB-cyan-pre-eGRASP (p30) were injected into BLA, VH and VTA, respectively. AAV2/1-CaMKIIα-iCre and AAV2/1-pAAV-EWB-DIO-myrTagRFP-T-P2A-post-eGRASP were injected into mPFC (Fig. 1A-C). All injections were performed unilaterally into the right VH, VTA, BLA and mPFC sequentially using 28-gauge microinjectors at a rate of 0.1 µL/min (coordinates from bregma are AP: -3.05, ML: -3.35, DV: -4.2 **for VH;** AP: -2.87, ML: -0.35, DV: -5.1 **for VTA;** AP: - 1.2, ML: -3.42, DV: -5.4 **for BLA;** and AP: +1.3, ML: -0.3, DV: -2.3 **for mPFC**). For the details of viral genome densities and volumes injected in each region please check Supplemental Table-1. Rectal temperature was monitored throughout the surgical procedure. Flunixin meglumine (2.5 mg/kg, sc) was administered for analgesia after the surgery.

**Figure 1:**
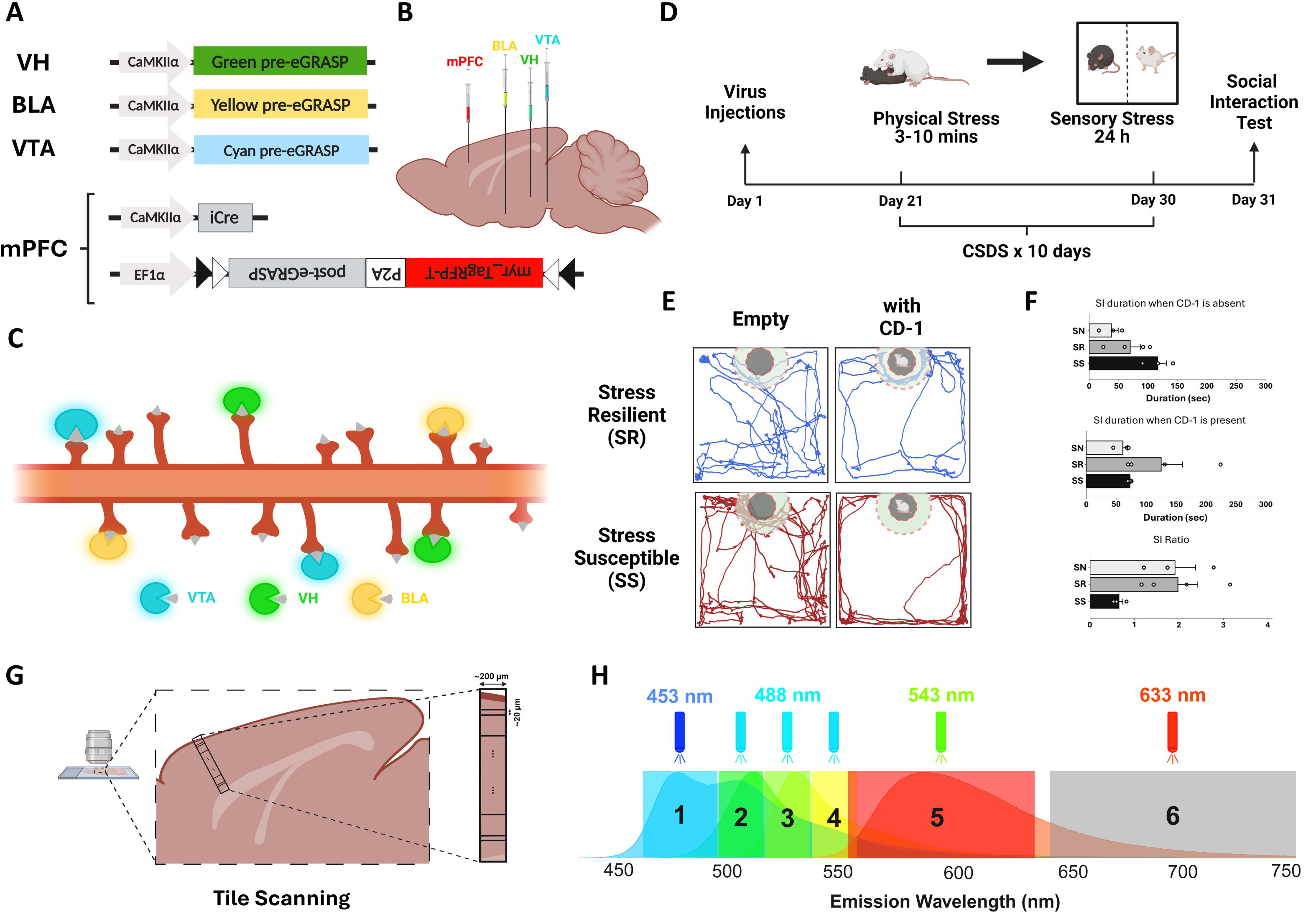
Description of the Experimental Strategy. **A:** Illustration of the pre- and post-synaptic AAV constructs. **B:** Illustration of the AAV injection sites, colors corresponding to specific e-GRASP constructs expressed by AAVs. **C:** The schema depicts a pyramidal mPFC neuron dendrite, tagged with TagRFP-T, illustrating synapses formed with afferents from BLA, VH and VTA, each labeled with a different color. **D:** Timeline of the experimental procedure. **E:** Representative tracking images during the SI test show SR (top) and SS (bottom) mice. **F:** SI durations in the absence and presence of a CD1 mouse, and SI ratios are presented as mean ± SEM. **G:** The schema of tiled imaging, covering all layers of a mPFC column. **H:** Excitation (top) and emission wavelengths (bottom) used for multispectral confocal imaging are specified. Numbers 1-6 indicate emission windows. Emission wavelength curves were taken from *fpbase.org*(*36*). Abbreviations SN: stress-naïve; SR: stress-resilient and SS: stress-susceptible.

### Chronic Social Defat Stress and Social Interaction Test

Three weeks after the injections, mice were exposed to 10-day chronic social defeat stress (CSDS) (Fig. 1D). Stress-naive (SN) control mice were kept under similar conditions, with their cage mates kept on the other side of the plexiglass barrier instead of the aggressor mice.

CSDS was carried out as Golden et al described (31). Briefly, an intruder C57BL/6 male mouse was placed in the cage of an aggressive CD-1 mouse. Duration of physical interaction was determined with an algorithm for each mouse based on the occurrence of bleeding to prevent injury. Then the cage was partitioned with a plexiglass barrier allowing visual, olfactory and vocal interactions and mice were cohoused for 24 hours. The stress protocol was repeated for 10 days with a different CD-1 mice. On day 11, stressed animals were divided into stress susceptible (SS) (n=3) and stress resilient (SR) (n=4) groups according to their social interaction (SI) ratios (Fig. 1E-F). Briefly, an interaction cage (9.5 cm in diameter) was placed at one edge of a 45 x 45 cm arena. All C57BL/6 mice were habituated to the arena for 3×20minutes. Then the time spent by the C57BL/6 mouse around the interaction zone (circular area with a diameter of 23.75 cm, sharing the same center with interaction cage) when the cage is empty and when it contains an unfamiliar novel CD-1 male mouse was recorded with Ethovision XT 8.0 (for 5 minutes/each). SI ratio was calculated by dividing the time spent around the interaction cage when CD-1 mouse was present to the time spent around the interaction cage when interaction cage was empty. Mice were classified as SR if the SI ratio is >1 and SS if the SI ratio is <1. SN mice (n=3) had SI ratio higher than 1, i.e. spent more time around the cage when a social target was present (Fig. 1F) (31).

### Tissue Preparation

Mice were perfused with heparin and 4% paraformaldehyde (PFA) under deep anesthesia. The brains were immersed in 4% PFA solution at +4°C overnight, followed by washing in cold phosphate-buffered saline (PBS) for 3 days to minimize PFA-induced autofluorescence. They were then immersed in a 30% sucrose solution for 2-5 days and stored at -80°C until sectioning. For sectioning, the frozen whole brain was placed on the tissue holder with the lateral part of contralateral hemisphere facing upward with 10-15 degrees oblique angle and was immersed in the embedding medium. Free floating 60 µm thick sagittal sections were obtained and mounted with 1/1 : glycerol/PBS solution.

### Imaging of Synapses and Dendrites

The images were taken from mPFC columns that clearly display dendritic signals, indicating efficient expression of myrTagRFP-T, and were collected by the same investigator who conducted the SI test. Multispectral, high resolution 3D images were acquired from mPFC across the full length of the cortical column (200 x 1500 x 30 μm^3^) by a Zeiss LSM810 laser-scanning confocal microscope, using a 40× oil objective (NA: 1.3) (Fig. 1G-H). The x, y pixel size and z-step size were determined to ensure optimal acquisition of the cyan eGRASP signal, which has the lowest emission wavelength among the signals. Accordingly, the pixel size and z-step size were adjusted to 0.103 µm and 0.3µm, respectively. Then, 30 µm-thick overlapping (10%) tile-scans with field of view (FOV) of 205 × 205 µm^2^ spanning the full cortical thickness from surface to corpus callosum were obtained. Then images from all tiles were stitched to form a full-length cortical column. For multispectral imaging, the same FOVs were imaged using 453, 488, 543 and 633 nm laser excitations. Emissions of fluorescent proteins were bandpass-filtered as follows: 465-490 nm for cyan eGRASP under 453 nm excitation; 495-515 nm, 515-535 nm, and 535-555 nm for green and yellow eGRASP under 488 nm excitation; 550-630 nm for TagRFP-T under 543 nm excitation; and 645-750 nm for autofluorescent signals under 633 nm excitation.

### Reflectance Imaging

Confocal reflectance imaging was performed to determine the boundary between layer VI and corpus callosum. Images were obtained in tile scans by a Leica SP8 confocal microscope. Briefly, a narrow band reflectance signal was obtained by excitation at 488 nm and emission at ∼483-493 nm using a RT15/85 beamsplitter. Z-stacks were obtained with 2-3 µm step size. The pixel size for this setting was ∼0.6 µm. Dendritic TagRFP-T labeled signals detected by 552 nm excitation were overlaid on the reflectance signal. Finally, images were stitched via the stitching algorithm of Leica Application Suite X (LASX) v3.5.5.19976.

### Image Processing

Image processing pipeline is shown in Figure 2 and includes stitching of the tiles, spectral unmixing of synaptic and dendritic signals deconvolution, and segmentation of the dendritic signals with an artificial neural network model (32) and reconstruction of dendritic segments (Fig. 3A-I, Suppl. Fig. 1, Suppl. Vid. 1-2).

**Figure 2:**
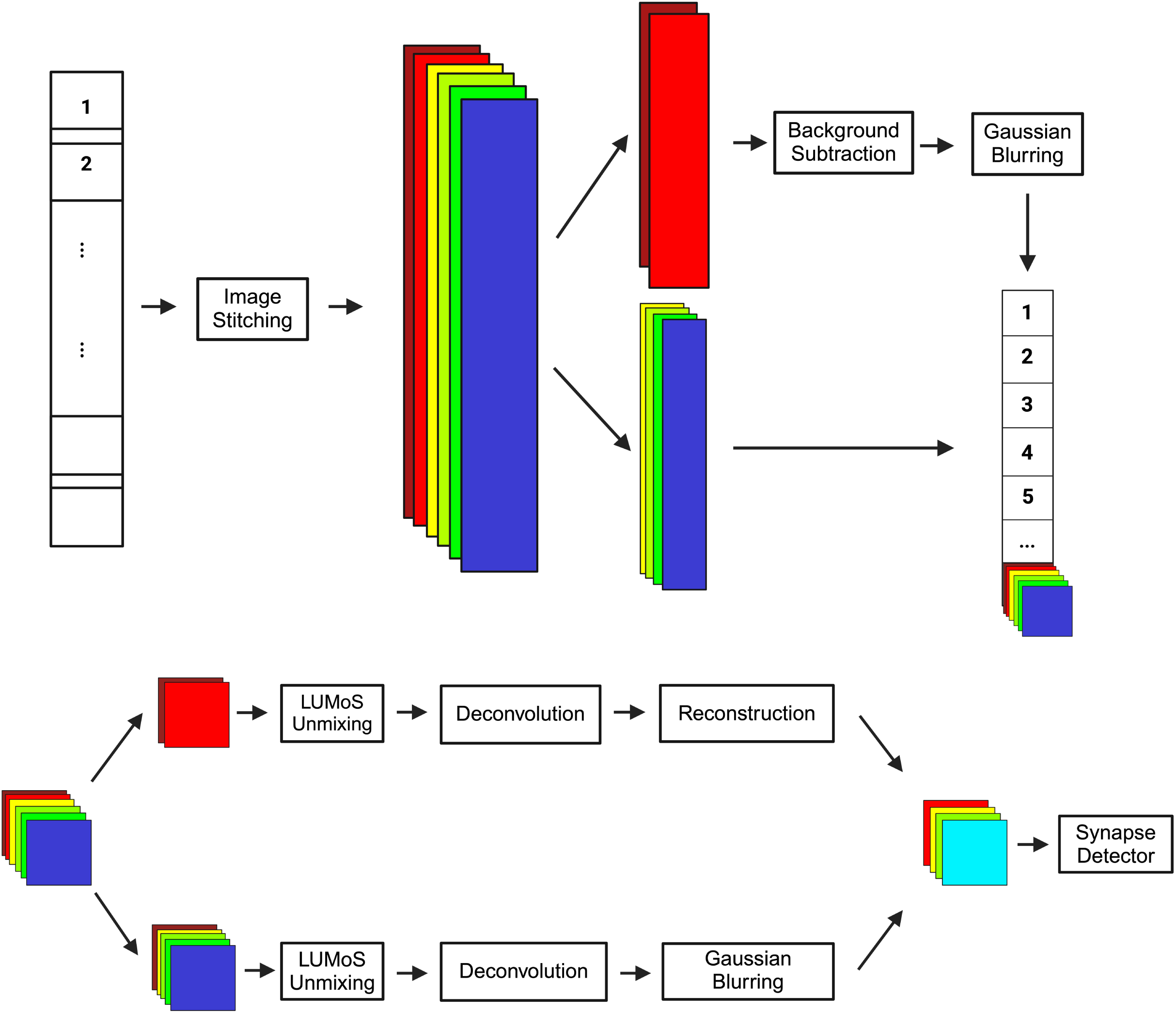
Image processing pipeline: Top: The framework for processing tiled images. Bottom: Framework for handling individual tiles: The dendritic image processing is illustrated at the top, while the processing of the synaptic images is depicted at the bottom.

**Figure 3:**
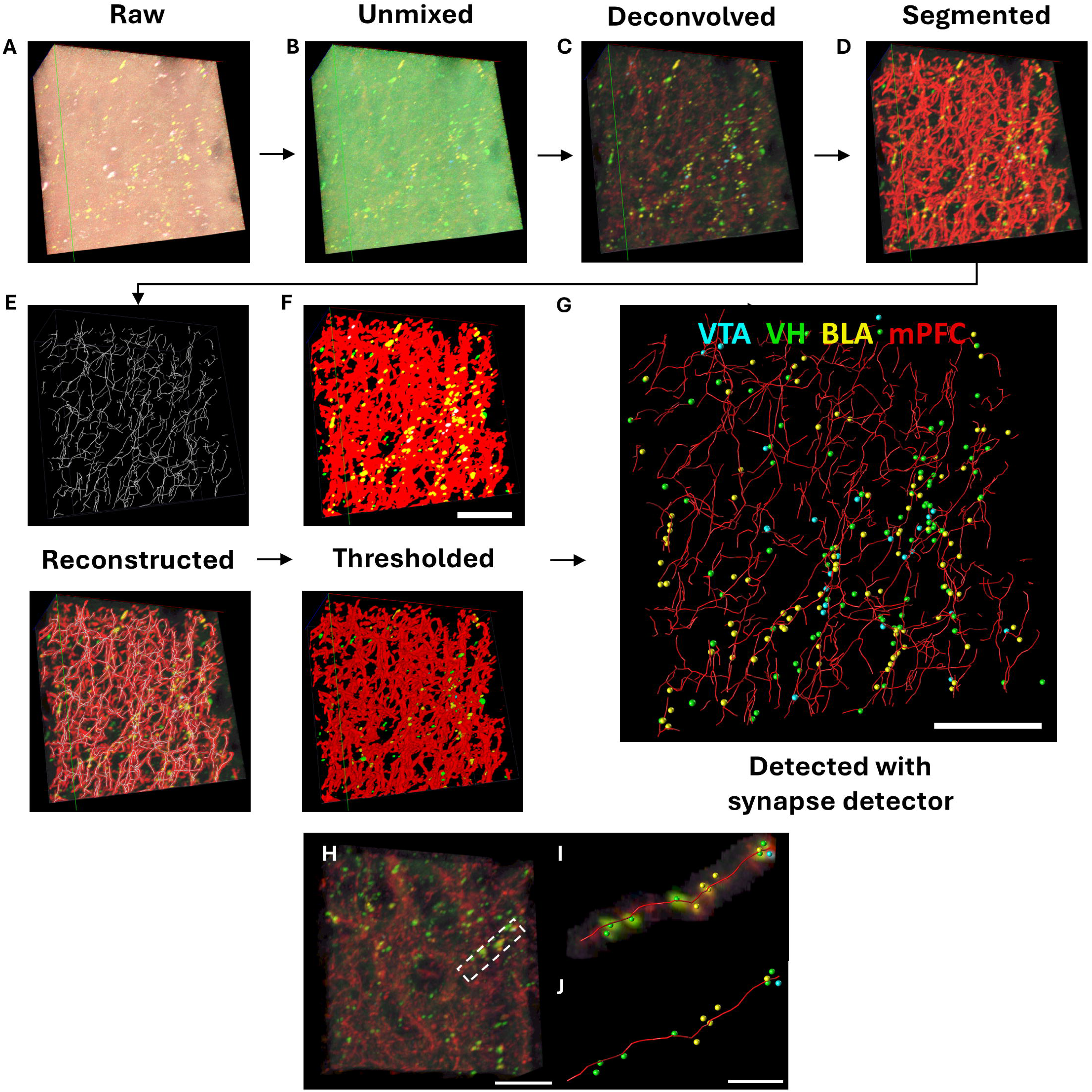
Processing of individual tiles and synaptic signals to visualize synapses between long-range inputs from the VH, BLA, and VTA, and dendrites of the mPFC. **A-F:** Representative images from different steps of the image processing pipeline. Image processing begins with **A:** the raw image and involves several steps: **B:** Unmixing, **C:** Deconvolution, **D:** Segmentation, **E:** Top: Dendritic reconstruction, Bottom: Segmented stack image overlaid on dendritic reconstruction, and **F:** Thresholding: Top: Thresholded image stack. Bottom: Alpha blended version of the thresholded image. **G-J:** Processing of the synaptic signals. **G:** Synapses detected by Synapse Detector (Vaa3D) on the reconstructed dendritic segments, ball-and-stick view. **H:** Deconvolved stack sample. **I-J:** Zoomed-in image of a sample dendritic segment within the white dotted line box in Figure 3H. **I:** Dendritic segment reconstruction overlaid on the deconvolved image. **J:** Ball-and-stick view of dendritic segment and its synapses. (Scale bars for figures: Fig. 3A-G: 25 μm, Fig. 3H: 20 µm, and for Fig. 3I-J: 5 µm).

#### Stitching

Acquired tiles were stitched together using the Zeiss Stitching algorithm (ZEN

3.2 Black Edition) or BigStitcher plug-in of FIJI (33). The correlation threshold was kept above “0” for Zeiss Stitching algorithm. For BigStitcher application x1 downscaling was applied to prevent loss or duplication of signals in the overlapping regions.

#### Spectral Unmixing

Due to the overlapping emission wavelengths of the signals, they were unmixed using the Learning Unsupervised Means of Spectra (LUMoS) algorithm (Fig. 3A-B, Suppl. Fig. 1) (34). The unmixing process was repeated at least 50 times with 100 iteration and the reference image was selected from tile images and axial slices, where all signal subgroups (from all fluorophores as well as nonspecific autofluorescence) were observable within the FOV. Unmixing of dendritic from autofluorescent signals and unmixing of synaptic signals (from each other and from autofluorescent signals) were done in two separate processes.

To validate LUMoS for unmixing cyan, green, yellow eGRASP signals from each other, spectral lambda scans were used (Suppl. Fig. 2). Lambda scans were acquired via Zeiss LSM810 laser scanning confocal microscope. Briefly, a ∼105 × 105 µm^2^ area was scanned with 1024 × 1024 pixels using a 40× oil objective (NA: 1.3). Excitation was performed with 453 and 488 nm lasers, to detect spectral emission profiles of cyan and green/yellow eGRASP, respectively. Fluorescence emission was collected in 5 nm-wide non-overlapping bands, between 460-575 nm for cyan and 495-600 nm for green/yellow eGRASP. Multispectral fluorescence data for comparison was collected at the same FOV prior to the lambda scans, as described above, and subsequently submitted to the LUMoS algorithm. Lambda stacks were further processed with Stowers Spectral Unmixing plug-in of FIJI (35), to extract the emission spectra of specific eGRASP proteins. In a region of interest with three different types of synaptic signals in the field, we first confirmed the presence of CFP, GFP and YFP, via checking their spectral emission profiles (Suppl. Fig. 2A-C) with reference spectra acquired from fpbase.org (36). Then, we processed the image data from the same field with LUMoS. We found ∼100 % overlap (n = 4 stacks) between the LUMoS classification of CFP, YFP and GFP and with their ground-truth synapse distribution, respectively (Suppl. Fig. 2D-E).

#### Deconvolution

To be able to generate a 3D reconstruction of the dendrites and synapses, deconvolution of the synaptic and dendritic signals was performed using GPU-based Richarson-Lucy algorithm (37). First, we prepared images by making the voxels isotropic via Reslice Z function in FIJI. Then point spread function (PSF) with the size of 256 x 256 x *N*, where *N* is the number of z-stacks of the tile image, was prepared by using Diffraction PSF 3D plug-in of FIJI. Optimum wavelengths of eGRASP proteins and TagRFP-T were used for PSF generation (28). Finally, Richardson-Lucy deconvolution algorithm with 20 iterations was applied for all signals (Fig. 3C). Synaptic signals were further processed by Gaussian filtering with a window size of 3 pixels. Stitched (BigStitcher) and unmixed (LUMoS) images were processed in Fiji. Synaptic and dendritic images were then merged according to their tile locations.

#### Reconstruction

We reconstructed the dendrites using the automated approach in Gliko et al (2024) that combines U-Net convolutional neural network segmentation with post-processing to produce a digital reconstruction (32). Many attempts on automating the segmentation of neuronal arbor morphology from optical sectioning microscopy exist in the literature (38–41). Gliko et al (2024) describes an automated reconstruction method based on training an artificial neural network on brightfield microscopy images of mouse cortical neurons (Fig. 3D,E). We used this method for automating reconstruction of dendritic segments in our dataset because (i) the low SNR of brightfield microscopy may lead to more effective suppression of spurious branch detection, and (ii) the voxel size of the dataset in Gliko et al (2024) and our images are similar. We note that training in Gliko et al (2024) was performed on “inverted” brightfield images so that the images resembled those from fluorescence microscopy (32). Segmentation was post-processed by thresholding the background, connected component analysis to remove short segments, skeletonization, and conversion to the .swc format, followed by pruning and down sampling. Finally, we quantified the accuracy of this approach by manually checking the 3D alignment of randomly selected 10 dendritic segments from each dendritic reconstruction (150 segments with total length of ∼1290 μm from ∼2000 x 700 x 300 pixel sized 119 images) with the deconvolved dendritic image, resulting in 96% accuracy (Suppl. Fig. 3).

We did not determine the diameters of the dendritic branches. We also did not classify the dendrites into apical and basal compartments since the directionality with respect to the soma could not be determined during imaging.

### Definition of Dendritic Segments

Dendritic signals across all mPFC layers labeled with TagRFP-T were intermingled, making it difficult to determine whether they belonged to the same dendritic tree. This challenge was due in part to the diffraction limit (0.25 - 0.35 µm) of the confocal microscopy, as well as to erroneous linkages between dendritic segments introduced during dendritic reconstruction. Therefore, we adopted a conservative approach by defining dendritic segments as portions of the reconstructed tree located between two adjacent branching or leaf nodes and evaluated synapses only on these short segments. Segments affected by image edge artifacts were excluded during quality control, and further quality control measures were applied to remove spurious segments with a length of 0 μm.

### Synapse Detection

BLA-mPFC, VH-mPFC and VTA-mPFC synapses were detected using the Synapse Detector plug-in of Vaa3D (42), which identifies synapses via adaptive watershed segmentation within a specified distance from the dendritic reconstruction (Fig. 3G-I). We ran this plug-in by the following five steps:

1. We thresholded deconvolved synaptic and segmented dendritic signals. Threshold values for synapses were determined by comparing raw synapse data and deconvolved synapse data (Fig. 3F, Suppl. Fig. 1). Threshold values for dendritic images were determined to avoid discontinuities along dendritic branches.
2. We prepared the 3D 8-bit RGB stacks that the Synapse Detector requires as input as follows:

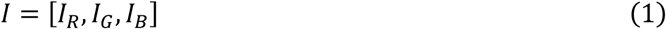

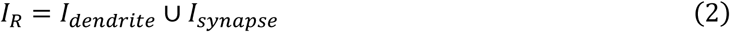

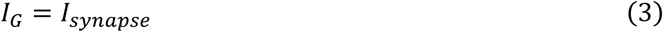

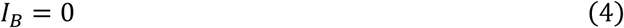

*I*_*R*_, *I*_*G*_and *I*_*B*_indicate 8-bit images, sized H x W x Z (height x width x depth of the image stack). *I*_*dendrite*_ indicates the segmented dendritic image stack, *I*_*synapse*_ indicates the synapse image of size H x W x Z. *I*_*R*_ is the union of dendritic (*I*_*dendrite*_) and synaptic image (*I*_*synapse*_) stacks. *I*_*B*_ is a stack of all zeros since the the algorithm writes the detected synapse information on this third channel after detection.
3. Synapse Detector has 4 main parameters: minimum signal density threshold values for dendritic and synaptic images, minimum synapse volume and maximum search distance from.swc tree. We did not use the signal density threshold values for dendritic and synaptic images as we applied thresholding in step 1. We limited the analysis to those synapses detected within ≤ 20 pixels (∼ 2 µm) distance around the swc tree (43) and those occupying at least 30 voxels (∼0.030 µm^3^) as the diffraction limit (0.25 - 0.35 µm) requires a detectable signal to occupy at least 3 pixels space (∼ 0.3 µm) along each dimension.
4. The Graphical User Interface (GUI) of the Synapse Detector allows the user to examine the synapses segment-by-segment, include/exclude detected synapses from the analysis, adjust the detected synapse volumes, and annotate synaptic positions as dendrite-targeting or spine-targeting. While human-annotated synaptic mapping is the gold standard way of input mapping, such manual inspection is infeasible for large datasets. Therefore, we visually examined the automatically detected synapses using only a zoomed-out view within the GUI of Synapse Detector. If most of the synapses have been tagged as detected, we have extracted the outputs of the algorithm in .csv format.
5. We repeated the above 4 steps for each of the three different colored synapse separately. The outputs of Synapse Detector used in this study are: **i)** 3D spatial locations of detected synapses, **ii)** Volumes of the detected synapses, and **iii)** For each synapse, the closest node to the detected synapse on the dendritic segment was identified, along with the corresponding dendritic segment and tree information.

### Comparison Between Automatic and Manual Counting Results

We checked the accuracy of the Synapse Detector in automatic synapse detection by two means:

1. In randomly selected 3D stacks (n = 49 images with a size of ∼600 x 2000 x 300 pixels from 15 cortical columns) from different layers, we confirmed each automatically detected synapse by visually inspecting and locating it in the unmixed images. Visual comparison was performed in Fiji by comparing detected synapses and the unmixed synaptic signals. Subsequently, the percentage of matched synapses among the total number of automatically detected synapses was calculated. This method suggests that 95.7% of automatically labeled synapses were truly synaptic signals and the percentage of true synapses was similar regardless of the brain region of origin of mPFC afferents (Suppl. Fig. 4A,B).
2. We manually counted all three types of synapses (formed by projections from BLA, VH and VTA) in randomly selected 3D frames from different layers on 3D visualization module of Vaa3D. Only signals within ∼2 µm from the reconstruction tree were manually counted, as Synapse Detector was set to detect synapses within ∼2 µm of a reconstructed dendrite. After manual labeling of synapses, spatial locations of the synapses were saved as a ”.marker” file. We then computationally matched the spatial locations of manually detected synapses with automatically detected synapses using the Hungarian algorithm (44). Based on this, we initially found that 80.64 ± 11.74% of the manually annotated synapses were matched to automatically detected synapses, and 23.97 ± 12.13% of the automatically detected synapses were not annotated by the expert. Upon closer inspection, however, we found that the mismatches were overwhelmingly due to synapses with small volumes, making them more likely to be missed by the human (Suppl. Fig. 4C). We also observed that majority of the unmatched manual annotations (∼11.4% of all manually labeled synapses) were assigned to incorrect locations by the graphical user interface of Vaa3D, which infers the 3D location from a click on the 2D screen. After exclusion of these mislocated manual annotations from the calculation and inclusion of the smaller synapses, we found that the overall synapse detection performance was ∼91%.

### Identification of mPFC Layers

Boundaries between cortical layers were identified by using the parcellation scheme of Common Coordinate Framework of Allen Brain Atlas (45). Accordingly, layer I, II/III, V and VI have been expected to cover 8%, 14%, 61% and 17% of the cortical column, respectively. Three dimensional cortical reconstructions were divided into laminae based on these ratios. Layer V was further divided into 2 equal halves: layer Va and layer Vb. The mean ± standard deviation of the thickness for each layer were 113.83 ± 21.46 µm (Layer I), 194.72 ± 37.14 µm (Layer II/III), 858.31 ± 154.35 µm (Layer V) and 239.20 ± 43.02 µm (Layer VI). Segments straddling two neighboring layers were excluded from the analysis.

### Calculation of Percentages of Each Synapse-to-all Detected Synapses

After dividing cortical columns into cortical layers, we calculated the ratios of synapses made by each long-range projection originated from a certain brain region (represented by color in the formula) to all detected synapses from all three brain regions. Let 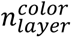 indicate number of synapses from BLA, VH or VTA in the layer I, II/III, Va, Vb or VI:

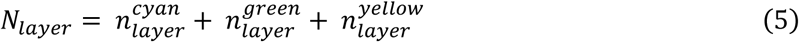

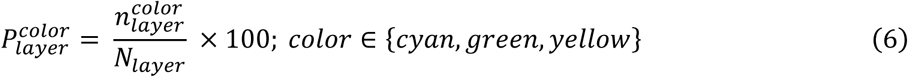

*N*_*layer*_ indicates numbers number of total detected synapses in a certain cortical layer. 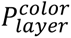 indicates the percentage of certain detected synapses color in given layer. We than created a color percentage matrix *M*^*color*^for each color:

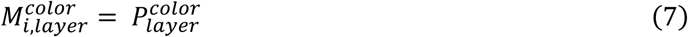

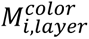 indicates the percentage value of cyan, green or yellow eGRASP labeled synapses in certain *layer* of cortical column *i* where *i* is the identity of cortical column. The expected size of the color percentage matrix is 5 x 13 (parts of cortical columns x number of mice) for each brain region projecting to mPFC.

### Dimensionality Reduction Matrix for Principal Component Analysis

To apply dimension reduction first, *M*^*color*^ matrices belonging different brain regions were concatenated in 2 dimensions.

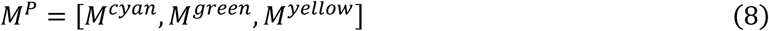

The final size of dimension reduction matrix (*M*^*P*^) was 13 x 15. Then, principal component analysis was performed. The first and second principal components have been used for plotting low dimensional projection of percentages of synapses made by single brain region-to-all detected synapses matrix.

### Formation and Analysis of Closest Spatial Distance Matrix of BLA and VH Synapses

For BLA and VH synapses in layer Vb and layer VI, the first four closest spatial distances to BLA or VH synapses were measured. A 42,317 x 8 sized matrix was formed by appending the closest VH and BLA synapse distances, respectively. Then Uniform Manifold Approximation and Projection (UMAP) was used to visualize synapses, behavioral profiles and SI ratios (46). Default values (version 0.5.5) were used for the hyperparameters of the UMAP code.

Random Forest Classifiers were formed using 10 forests with 75 trees. K-fold cross validation scheme with 10 independent parts was applied to each forest. In each fold, 10% of the dataset has been used as testing dataset. Only data labels have been randomized to train RF classifier to compare RF classification performance for structured and randomized datasets. Parametric measures of accuracy and F1 scores of RF classifier have been used for comparison between main data and randomly shuffled data. RF classifiers have been trained using sklearn v1.4.0 inside Python 3.9.

### Statistical Analysis

Sample size was determined based on similar studies in the literature (26, 47, 48). Data are presented as median with interquartile ranges, mean ± S.E.M or mean with 95% confidence interval. Kruskal-Wallis (KW) test was used for multiple group comparisons and followed by pairwise comparisons by Mann Whitney U (MWU) tests, p values for MWU were adjusted by Bonferroni correction. All analysis were two tailed. p value less than 0.05 and 0.0167 were considered as significant in KW and post-hoc MWU tests, respectively. All statistical analysis was performed using Scipy v1.11.1. Plots were drawn by using Matplotlib v3.8.0 and Seaborn v0.12.2.

### Code Availability

All the codes used are available from the corresponding author upon request.

## RESULTS

### Long-range inputs from VH, BLA and VTA to mPFC pyramidal neurons across the cortical column

The median total numbers of detected synapses were 2,501 (IQR = 1,077 – 6,293; n = 5 cortical columns from 3 mice); 7,509 (IQR = 2,560 – 9,177; n = 5 cortical columns from 4 mice) and 5,845 (IQR = 789 – 8,598; n = 5 cortical columns from 3 mice) in SN, SR and SS mice, respectively. In the SN group, VH-mPFC synapses constituted the highest fraction of all identified synapses (70.33 ± 16.82%) followed by BLA-mPFC synapses (16.09 ± 11.25%) and VTA-mPFC synapses (13.57 ± 6.09%). In the SR mice, on the other hand, fractions of VH-mPFC and BLA-mPFC were similar (49.62 ± 17.80% and 38.17 ± 20.37%, respectively; p = 0.69, MWU) followed by VTA-mPFC synapses (12.19 ± 4.32%). The percentages of BLA-mPFC, VH-mPFC and VTA-mPFC synapses in SR mice were similar to those observed in the SN group (p = 0.09, p = 0.15 and p = 0.69, MWU, respectively). In the SS group, the percentage of BLA-mPFC synapses tend to be higher than that of VH-mPFC synapses (54.27 ± 17.18% vs. 25.74 ± 15.12%, respectively; p = 0.055, MWU). The percentage of VTA-mPFC synapses in SS mice was comparable to that observed in the other two groups (19.97 ± 9.79%; p = 0.30, MWU).

### Layer and stress phenotype-dependent changes to the ratio of BLA synapses

We then examined the distribution of synapses across each cortical layer. Previous literature showed that BLA and VH axonal projections to the mPFC preferentially target layer II/III and layer V, respectively (8, 21–23). In line with the previous literature, we found that the percentage of BLA-to-all detected synapses (BLA/(BLA+VH+VTA)*100) was higher in layer II/III than in layers V and VI in SN group (p = 0.0079, p = 0.035, respectively, MWU test). Chronic stress exposure increased the percentage of BLA synapses in layer II/III compared to SN controls (Suppl. Fig. 5A,B). The percentage of BLA synapses in layer II/III was similar in SS and SR groups (Fig. 4A-C). On the other hand, the percentage of BLA synapses in layer Vb and VI increased only in the SS group and was significantly higher than those in the SN and SR groups (Fig. 4D,E). The percentages of BLA synapses in layer Vb and VI showed a significant negative correlation with SI ratio (Fig. 4F-H).

**Figure 4:**
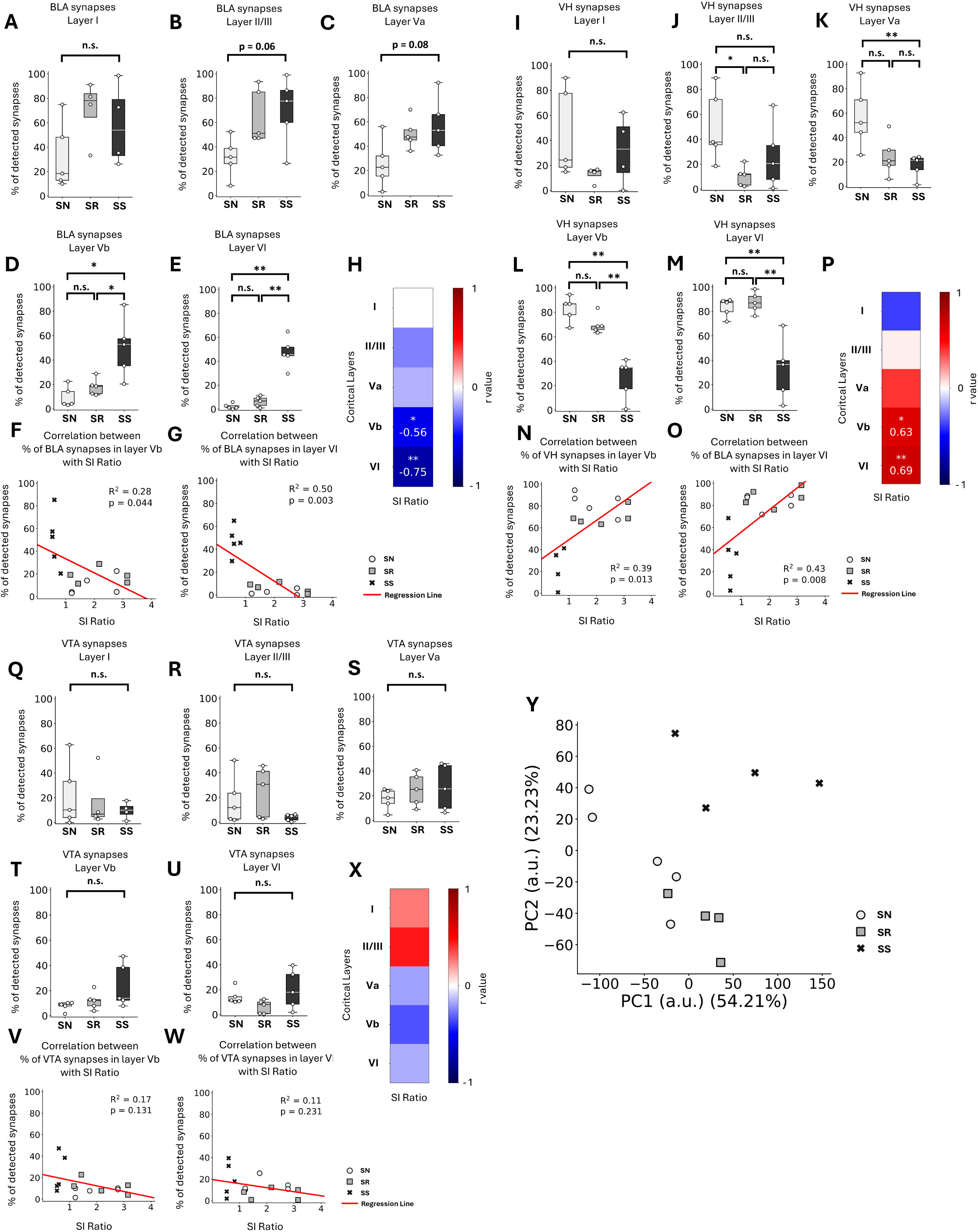
Layer- and stress phenotype-dependent changes of BLA, VH and VTA synapses. **A-E:** Percentage of BLA-to-all detected synapses in mPFC layers I-VI. **A:** Layer I (Kruskal-Wallis (KW) test: H = 4.02, p = 0.13); **B:** Layer II/III (KW test: H = 5.53, p = 0.06); **C:** Layer Va (KW test: H = 4.87, p = 0.08); **D:** Layer Vb (KW test: H = 8.78, p = 0.012; Mann-Whitney U (MWU) test: p = 0.0158, SS vs SR ; p = 0.0158, SS vs SN ; p = 0.22, SR vs SN); **E:** Layer VI (KW test: H = 10.50, p = 0.0052; MWU test: p = 0.0079, SS vs SR ; p = 0.0079, SS vs SN ; p = 0.15, SR vs SN). **F:** Correlation between the BLA percentages and the SI ratio in layer Vb (Spearman correlation: rho = - 0.56, p = 0.030; linear regression: F(1,13) = 4.96, R^2^ = 0.28, p = 0.044). **G:** Correlation between the BLA percentages and the SI ratio in layer VI (Spearman correlation: rho = - 0.75, p = 0.001; linear regression: F(1,13) = 12.97, R^2^ = 0.50, p = 0.003). **H:** Spearman correlations between BLA percentages and the SI ratio in each layer. Correlation coefficients of significant correlations were indicated on the linear heatmap. **I-M:** Percentage of VH-to-all detected synapses in mPFC layers I-VI. **I:** Layer I (KW test: H = 3.75, p = 0.15); **J:** Layer II/III (KW test: H = 6.02, p = 0.049; MWU test: p = 0.54, SS vs SR ; p = 0.15, SS vs SN ; p = 0.0158, SR vs SN); **K:** Layer Va (KW test: H = 7.57, p = 0.022; MWU test: p = 0.69, SS vs SR ; p = 0.0079, SS vs SN ; p = 0.055, SR vs SN, MWU test); **L:** Layer Vb (KW test: H = 10.82, p = 0.0044; MWU test: p = 0.0079, SS vs SR ; p = 0.0079, SS vs SN ; p = 0.09, SR vs SN); **M:** Layer VI (KW test: H = 9.50, p = 0.0086; MWU test: p = 0.0079, SS vs SR ; p = 0.0079, SS vs SN ; p = 0.69, SR vs SN). **N:** Correlation between the VH percentages and the SI ratio in layer Vb (Spearman correlation: rho = 0.63, p = 0.012; Linear regression: F(1,13) = 8.33, R^2^ = 0.39, p = 0.013). **O:** Correlation between the VH percentages and the SI ratio in layer VI (Spearman correlation: rho = 0.69, p = 0.0040; Linear regression: F(1,13) = 9.80, R^2^ = 0.43, p = 0.008). **P:** Spearman correlations between the VH percentages and the SI ratio in each layer. Correlation coefficients of significant correlations were indicated on the linear heatmap. **Q-U:** Percentage of VTA-to-all detected synapses in mPFC layers I-VI. **Q:** Layer I (KW test: H = 0.11, p = 0.94); **R:** Layer II/III (KW test: H = 2.47, p = 0.28); **S:** Layer Va (KW test: H = 1.26, p = 0.53); **T:** Layer Vb (KW test: H = 4.5, p = 0.10); **U:** Layer VI (KW test: H = 4.34, p = 0.11). **V:** Correlation between the VTA percentages and the SI ratio in layer Vb (Spearman correlation: rho = -0.34, p = 0.21; Linear regression: F(1,13) = 2.60, R^2^ = 0.17, p = 0.13,). **W:** Correlation between the VTA percentages and the SI ratio in layer VI (Spearman correlation: rho = - 0.16, p = 0.56; Linear regression: F(1,13) = 1.58, R^2^ = 0.11, p = 0.23). **X:** Spearman correlations between the VTA percentages and the SI ratio in each layer. Correlation coefficients of significant correlations were indicated on the linear heatmap. **Y:** Principal component analysis: Samples (synapse ratios per layer and afferent) plotted in two dimensions using their projections onto the first 2PCs and labeled according to stress exposure and susceptibility status. Data is presented as median and IQR for boxplots graphs. For layer I, n = 5 sections from 3 SN mice, n = 4 sections from 4 SR mice and n = 4 sections from 4 SS mice; for layers from II/III to VI, n = 5 sections from 3 SN mice, n = 5 sections from 4 SR mice and n = 5 secitons from 3 SS mice. Asterisks in the figures indicate ∗∗: p ≤ 0.01, and ∗: p < 0.05. Abbreviations SN: stress-naïve; SR: stress-resilient and SS: stress-susceptible.

### Layer and stress phenotype-dependent changes to the ratio of VH synapses

Changes observed in VH-mPFC synapses were similar to those seen in BLA-mPFC synapses, but in the opposite direction. Mice exposed to CSDS showed a decrease in the percentage of VH-to-all synapses [VH/(BLA+VH+VTA)*100] in layer II/III compared to SN controls (Suppl. Fig. 5A,C). We did not observe any differences in the percentage of VH synapses between SS and SR groups (Fig. 4I-K). In contrast, only SS mice showed a significantly decreased percentage of VH synapses in layer Vb and VI (Fig. 4L-M). The percentages of VH synapses in layer Vb and VI were significantly correlated with SI ratio (Fig. 4N-P).

### Invariance of the ratio of VTA synapses to the stress phenotype

The VTA-to-all synapses percentage (VTA/(BLA+VH+VTA)*100), was similar across all layers of the mPFC in all groups (Fig. 4Q-U, Suppl. Fig. 5A,D). We did not find statistically significant correlations between SI ratio and the percentages of VTA synapses across different layers (Fig. 4V-X).

### A low dimensional representation of the relative abundances of VH, BLA and VTA synapses across the cortical column visually differentiates SS mice from SR mice

To further examine the dissociation of groups according to their stress susceptibility, we examined the low-dimensional structure of synaptic distributions across all layers using principal component analysis (PCA) in 13 cortical columns with an intact layer I (2 columns lacking layer I were excluded from the analysis, due to damage resulting in 13 columns, presumably from the AAV injections). We converted the 3×13×5 (synapse type count x column count x layer count) synapse percentage matrix into a 13×15 matrix by arranging the synaptic percentage data from VTA, VH and BLA from each layer into a single row. We then applied PCA to this matrix and visualized the distribution of cortical columns in the 2-dimensional plane based on the top two principal components (PCs). Approximately 77.44% of the variance in the synaptic percentages across the cortical column was captured by these two PCs. Notably, the cortical columns associated with stress susceptibility occupied distinct locations in this low dimensional space (Fig. 4Y). This supports the finding that relative synaptic abundances of BLA, VH and VTA across the mPFC laminae can explain the behavioral differences between SR and SS mice.

### The abundance of dendritic segments receiving input from multiple brain regions increases in SS mice

To gain deeper insight into the effects of stress and the differences between SS mice and the other groups, we investigated the percentage of reconstructed dendritic segments that had two or more synapses in all layers (Fig. 5, Suppl. Fig. 6, 7). In layer II/III, all stressed mice, regardless of their susceptibility, showed a significantly higher percentage of dendritic segments receiving multiple (>1) inputs exclusively from the BLA compared to the SN group (Suppl. Fig. 6). On the other hand, in layer Vb and layer VI, the percentage of dendritic segments receiving more than one inputs from only VH was significantly lower and those receiving more than one inputs from only BLA tended to be higher in the SS mice than the other two groups. There was no difference in the percentage of dendritic segments receiving multiple inputs from only VTA among groups (Suppl. Fig. 7).

**Figure 5:**
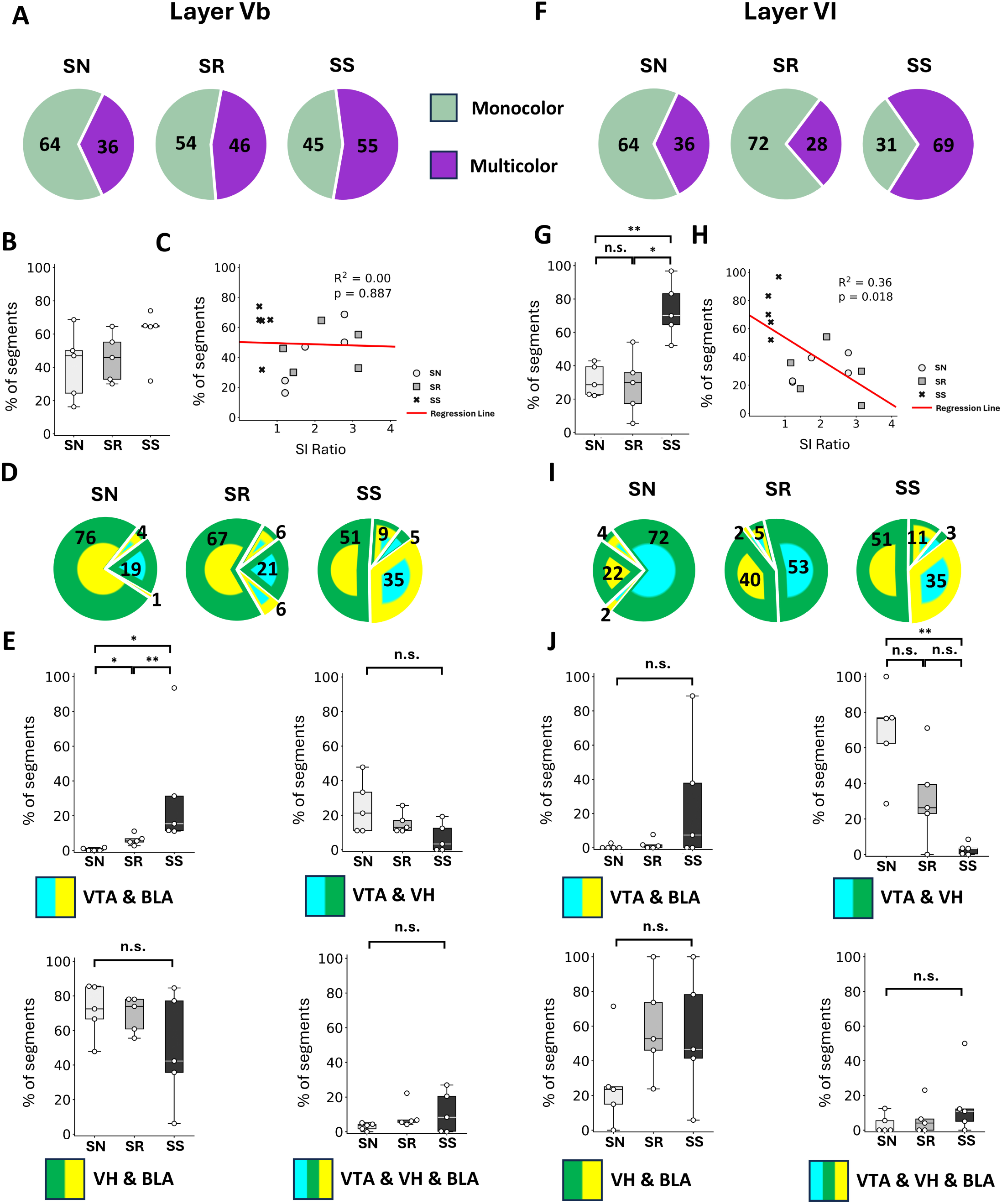
The number of segments forming synapses with multiple brain regions increased in Layer VI of SS mice. **A-E:** Analysis of dendritic segments receiving inputs from a single brain region (monocolor) and from multiple brain regions (multicolor) in layer Vb. **A:** Pooled data from each group showing percentages of dendritic segments receiving inputs from a single brain region (monocolor) and from multiple brain regions (multicolor) in layer Vb. The number of monocolor segments in SN, SR, SS mice were 1,082; 1,568 and 1,020, respectively. The number of multicolor segments in SN, SR, SS mice were 604; 1,317 and 1,237, respectively. **B:** Percentage of dendritic segments receiving input from at least 2 different brain regions in layer Vb (Kruskal-Wallis (KW) test: H = 2.96, p = 0.22). **C:** Correlation of the ratio of multicolor dendritic segments with the SI ratio in layer Vb (Spearman correlation: rho = - 0.18, p = 0.49; linear regression: F(1,13) = 0.02, R^2^ = 0.0016, p = 0.88). **D:** Pooled data from each group showing percentages of dendritic segments receiving input from multiple brain areas in layer Vb. **E:** Comparisons of percentages of dendritic segments receiving input from VTA & BLA *(KW test: H = 12.58, p = 0.0018; Mann Whitney-U (MWU) test: p= 0.0079, SS vs SR; p = 0.011, SS vs SN; p = 0.011, SR vs SN)*, VTA & VH *(KW test: H = 3.88, p = 0.14),* VH & BLA *(KW test: H = 1.93, p = 0.37)* and VTA & VH & BLA *(KW test: H = 3.31, p = 0.19).* **F-J:** Analysis of dendritic segments receiving inputs from a single brain region (monocolor) and from multiple brain regions (multicolor) in layer VI. **F:** Pooled data from each group showing percentages of dendritic segments receiving inputs from a single brain region (monocolor) and from multiple brain regions (multicolor) in layer VI. The number of monocolor segments in SN, SR, SS mice were 352; 1,589 and 465, respectively, while the number of multicolor segments in SN, SR, SS mice were 197, 628 and 1,015, respectively. **G:** Percentage of dendritic segments receiving input from at least 2 different brain regions in layer VI (KW test: H = 8.66, p = 0.013; MWU test: p = 0.015, SS vs SR; p = 0.0079, SS vs SN; p = 0.84, SN vs SS). **H:** Correlation of the ratio of multicolor dendritic segments with SI ratio in layer VI (Spearman correlation: rho = - 0.63, p = 0.010; linear regression: F(1,13) = 7.31, R^2^ = 0.36, p = 0.018). **I:** Pooled data from each group showing percentages of dendritic segments receiving input from multiple brain areas in layer VI. **J:** Comparisons of percentages of dendritic segments receiving input from VTA & BLA *(KW test: H = 2.61, p = 0.27)*; VTA & VH *(KW test: H = 9.08, p = 0.010; MWU test: p = 0.11, SS vs SR ; p = 0.0079, SS vs SN ; p = 0.055, SR vs SN)*; VH & BLA *(KW test: H = 3.46, p = 0.17);* VTA & VH & BLA *(KW test: H = 3.31, p = 0.19)*. Data is presented as median and IQR. Asterisks in the figures stand for ∗∗: p ≤ 0.01, and ∗: p < 0.05. For both layers, n = 5 from 3 SN mice, n = 5 from 4 SR mice and n = 5 from 3 SS mice. Abbreviations SN: stress-naïve; SR: stress-resilient and SS: stress-susceptible.

In both layer II/III and layer Vb, the percentages of dendritic segments receiving inputs from multiple brain regions were similar between groups (Suppl. Fig. 6, Fig. 5A-C). However, when layer Vb was further analyzed for each pair of multi-brain region originated synapses on individual dendritic segments, we found that the percentage of BLA&VTA synapses increased in both stressed groups, with a significantly greater rise in the SS group compared to the SR group (Fig. 5D-E). No such differences were observed between any synaptic pairs in layer II/III (Suppl. Fig. 6). In layer VI, on the other hand, the percentage of dendritic segments receiving inputs from multiple brain regions was significantly higher in the SS group than the other two groups (Fig. 5F-H). When analyzed separately for each pair, we observed a reduction in the percentages of VTA&VH synapses on the same dendritic segment in the SS group (Fig. 5I-J). Interestingly, we did not observe any difference in the percentage of BLA&VH synapses on the same dendritic segment among groups in either layer Vb or layer VI (Fig. 5E, 5J).

These findings suggest that the organism may adapt to stress by adjusting the number of same-brain region originated synapses on a single dendritic segment in mPFC deep layers. In contrast, the increase in multi-origin-synaptic pairs on the same dendritic segment observed in SS mice may signal a disorganization affecting the outputs of deep mPFC layers, possibly contributing to a dysfunctional stress response.

### The spatial organization of VH and BLA synapses on mPFC deep layers is disrupted in SS mice

To explore whether there is a general pattern of mPFC innervation by the VH and BLA on a large scale, we measured the Euclidean distances between VH and BLA synapses and determined the closest VH and BLA synapses for each VH and BLA synapse within the reconstructed mPFC volumes of ∼200 x 200 x 30 µm^3^ in layer II/III and ∼200 x 700 x 30 μm^3^ in layer Vb-VI. In layer II/III, both stressed groups—regardless of susceptibility or resilience—showed a tendency toward reduced distances between VH synapses and their closest BLA synapses (Suppl Fig. 8). All other distance measurements were similar among groups (Suppl. Fig. 8). In layer Vb and layer VI, there were no differences observed in the distances between VH synapses and their closest VH synapses, nor in the distances between BLA synapses and their closest BLA synapses across the three groups studied (Fig. 6A-B). However, we observed differential spatial organization of VH and BLA synapses relative to each other in SN/SR mice vs. SS mice: In the SN and SR mice, the distances of VH synapses to their corresponding closest BLA synapses were distributed relatively uniformly across a wide range (Fig. 6C), whereas the BLA synapses were preferentially located in close proximity (within 5 um) to VH synapses (Fig. 6D). In contrast, in the SS group, we observed a statistically significant decrease in the distances of VH synapses to their corresponding closest BLA synapses and a statistically significant increase in the distances of BLA synapses to their corresponding closest VH synapses (Fig. 6C-D).

**Figure 6:**
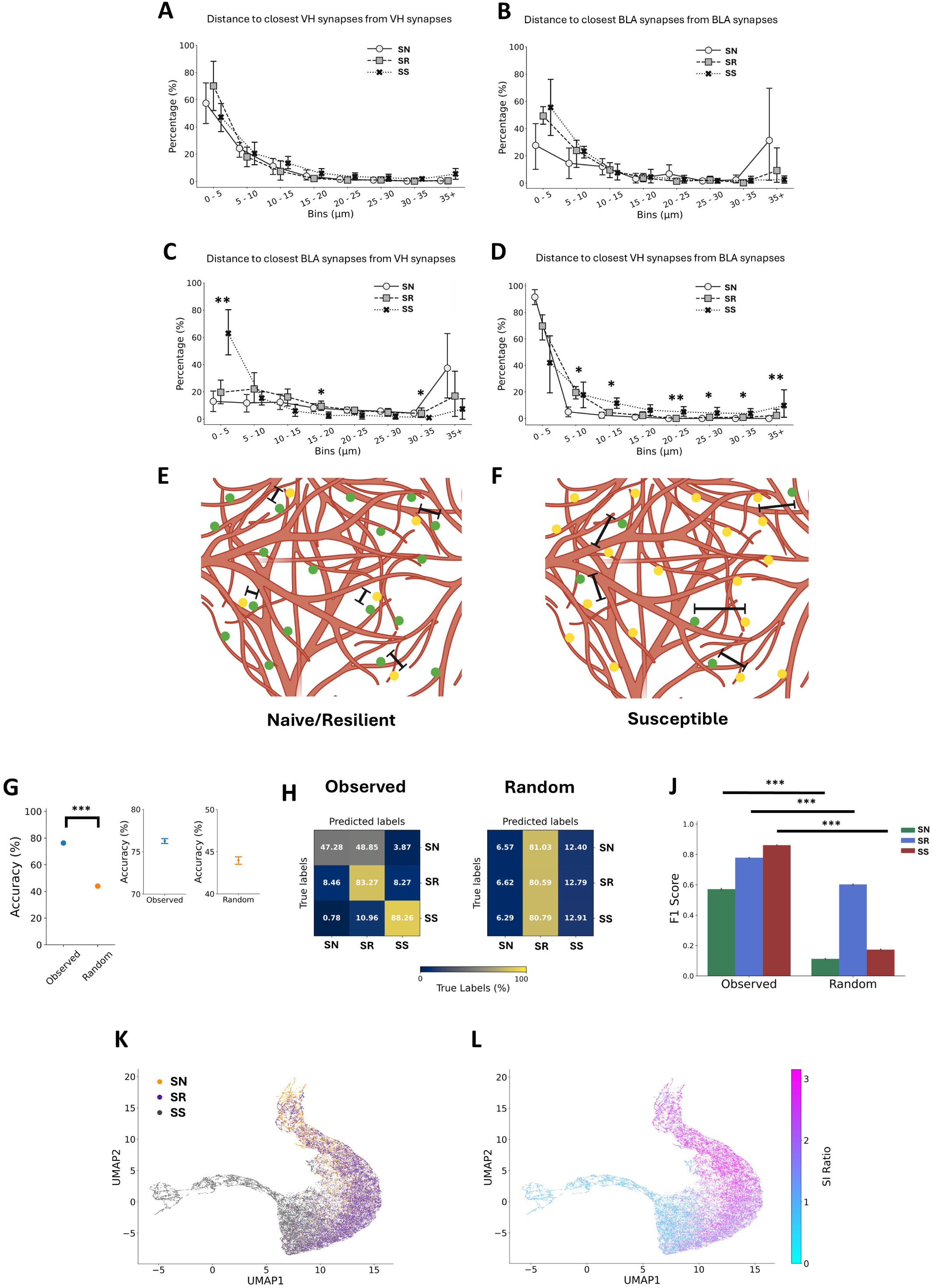
Euclidean Distances Between Unidentical, but not Identical Synapses were Distinct in SS mice. **A:** Histograms of Euclidean distances between each VH synapse and its closest VH synapse for different groups. **B:** Histograms of Euclidean distances between each BLA synapse and its closest BLA synapse for different groups. **C:** Histograms of Euclidean distances between each VH synapse and its closest BLA synapse for different groups. **D:** Histograms of Euclidean distances between each BLA synapse and its closest VH synapse for different groups. Percentages of each distance range was compared by Kruskal Wallis test for three groups, pairwise comparisons were carried out by Mann Whitney U test with Bonferroni correction (please see Suppl. Table 2 for detailed statistical values). **E-F:** Cartoon illustrations depicting the spatial organization of BLA and VH synapses in layers Vb and VI, highlighting the differences between SS mice and SN/SR mice. Panel E represents SN/SR mice, while Panel F shows SS mice. **G:** Comparison of accuracy after Random Forest Classification between observed and randomized data (p = 5.16 × 10^-5^, independent samples t-test). **H:** Confusion matrices of observed data (left) and randomized data (right). Values were normalized for the number of true labels. **J:** F1 scores for observed and randomized data for each behavioral group. For SN mice: p = 1.83 × 10^-24^; for SR mice: p = 2.34 × 10^-23^; for SS mice: p = 2.12 × 10^-30^, independent samples t-test. **K-L:** 2D UMAP projection of Euclidean distances to four closest VH and four closest BLA synapses. Panel K shows UMAP results colored by behavioral groups, Panel L shows colored by SI ratio. Data is presented as mean and 95% CI. ***: p < 0.001; **: p < 0.01; *: p < 0.05. n = 5 from 3 SN mice, n = 5 from 4 SR mice and n = 5 from 3 SS mice. Abbreviations SN: stress-naïve; SR: stress-resilient and SS: stress-susceptible.

These results suggest that the long-range projections follow structured, differential innervation patterns and are not random (Fig. 6E-F). We quantified this observation by training a random forest classifier (RFC) on a 42,317 x 8 matrix of closest synaptic distances, where each row corresponds to a synapse. In each row, the first four columns denote the Euclidean distances of that synapse to its four closest VH synapses, and the last four columns denote the distances to its four closest BLA synapses. This RFC had 76% accuracy in differentiating all three groups (Fig. 6G). Examining the confusion matrix reveals that the RFC can predict the SS group with ∼88% accuracy and the SR group with ∼83% accuracy. However, the prediction accuracy of the SN group is low (∼47%) (Fig. 6H). We observed that the classification accuracy and the F1 score (harmonic mean of precision and recall) decreases significantly when the RFC is trained on the same dataset with shuffled labels, simulating a scenario where synapses from the different regions are randomly distributed in the mPFC over the observed synaptic locations (Fig. 6G-J). Finally, we visualized this result using uniform manifold approximation and projection (UMAP) (46), a non-linear dimensionality reduction method: the synapses associated with the SR and SS phenotypes occupy different regions of the two-dimensional UMAP map of the same data matrix (Fig. 6K-L).

## DISCUSSION

While the stress response is a fundamental component of the behavioral repertoire of animals, a mechanistic, circuit-level understanding of it in health and disease remains incomplete. This is, in part, due to the complexity of the distributed circuitry that governs the behavioral responses, involving numerous synaptic connections between multiple brain regions. Here, by using multiple eGRASP constructs, multispectral volumetric imaging, computational tools for dendrite and synapse identification, and statistical methods for analyzing the resulting large-scale dataset, we aimed to elucidate this circuitry in the mouse. To our knowledge, this is the first study that mapped VH-mPFC, VTA-mPFC and BLA-mPFC synaptic connections separately across all cortical layers and compared them between SN, SS and SR mice. This multi-pronged approach revealed distinct patterns of organization of synaptic inputs to deep and superficial mPFC layers in the stress response and susceptibility.

Previous literature showed that BLA projections preferentially target layer II (21, 22, 49), whereas, VH projections are largely restricted to layer V of mPFC (22, 23). Consistent with these findings, we observed a similar organization of long-range synaptic inputs to the cortical layers of SN mice. CSDS altered the ratios of BLA-to-all synaptic inputs [BLA/(BLA+VH+VTA) and VH-to-all synaptic inputs [VH/(BLA+VH+VTA)] in mPFC superficial layers, regardless of the mice’s stress susceptibility. This indicates that changes observed in these layers are related to stress exposure itself, rather than to stress susceptibility. Previous studies reported reductions in dendritic length and spine numbers in mPFC layer II/III after chronic stress exposure (1–5). Our results, when considered together with these findings, suggest that mPFC neurons that receive input from VH were preferentially atrophied by chronic stress resulting in a decrease in VH inputs, whereas those that receive input from BLA were unaffected. Supporting this suggestion, Shansky et al (2009) demonstrated that pyramidal neurons projecting to entorhinal cortex showed dendritic atrophy in response to chronic stress, whereas those projecting to amygdala were resilient to stress-induced dendritic remodeling (7). Considering that there is a reciprocal connection between BLA and mPFC (i.e. BLA inputs preferentially innervate BLA-projecting mPFC neurons), one may speculate that BLA input receiving mPFC neurons are also resilient to atrophy (21, 22, 49, 50).

The organization of long-range inputs to deep layers of mPFC was disrupted only in the SS group. Not only did BLA-mPFC synapses predominate over VH-mPFC synapses, but the distance between BLA-mPFC synapses and their closest VH-mPFC synapses also increased in layer Vb and VI. Brain networks connecting mPFC, BLA, and hippocampus are involved in the regulation of fear-related and social behaviors (14, 17, 51–54). The interplay between these three regions in the control of anxiogenic/social behaviors is complex and may both promote and suppress fear or social behavior (17, 22, 24, 50, 55). Inputs from BLA and VH to the mPFC have been reported to have opposing effects on these behaviors (14, 17, 56). Intriguingly, within each of these brain regions, distinct populations of neurons with distinct connections encode “safe” and “aversive” cues, enabling the organism to react optimally to various stimuli (24, 25, 50). Our findings also revealed a delicate balance in the relative numbers and spatial organization of BLA and VH inputs in deep mPFC layers, which, when disrupted, may contribute to susceptibility to stress-related mental disorders. Selective activation of distinct neuronal ensembles within the mPFC may serve as a mechanism to switch between alternative behavioral responses by integrating sensory, contextual and emotional information, fine-tuning the expression and suppression of fear-related and social responses (24, 55, 57). Our findings of disrupted relative abundances and spatial organization of VH and BLA synapses, along with multi-brain region-originated synapses on single dendritic segments in SS mice, suggest that these alterations may interfere with signaling within the associated neuronal circuits and potentially hinder an adaptive stress response.

Mice are social animals and they tend to spend more time with social targets (stranger mouse) compared to non-social targets (i.e. objects) (58). Following CSDS, SS and SR mice demonstrate opposite behavioral responses when encountering a novel mouse in a novel context: SS mice exhibit social avoidance, while SR mice demonstrate social approach. This suggests that SR mice can discriminate threat-related stimuli or contexts from novel stimuli or contexts, whereas SS mice tend to overgeneralize the threat to all novel stimuli or contexts they encounter. Overgeneralization of fear is a characteristic feature of anxiety disorders and thought to be one of the mechanisms underlying posttraumatic stress disorder, panic disorder and generalized anxiety disorder (59–62). The increase in the relative abundance of long-range inputs from the BLA and the decrease in those from the VH to the deep layers of the mPFC pyramidal neurons in SS mice, as observed in the current study, could potentially impair the discrimination between threat-associated and non-threat associated stimuli. This disruption may then compromise the mPFC’s ability to produce an adaptive response. Intriguingly, a previous study showed that the discrimination of aversive cues from safe cues was associated with synchrony between mPFC and BLA activities in the theta frequency range (55). The entrainment of BLA activity by mPFC activity in the theta frequency range was reported in only those mice that can discriminate conditioned auditory tones from unconditioned tones (55). Additionally, the VH also synchronizes with mPFC activity at the theta frequency range during the exploration of anxiogenic novel environments, facilitating the discrimination of safe and aversive components of the environment (18, 19, 57). VH is also implicated in the formation and storage of social memory (53). VH neurons are more strongly activated in response to a familiar mouse compared to a stranger mouse, and their optogenetic inhibition interferes with the discrimination of familiar and unfamiliar mice (53). In parallel with these findings, our results suggest that the balance between BLA and VH inputs to the mPFC is crucial in associating social cues with aversion, and in discriminating safe cues from aversive cues to produce an adaptive response (Please see Fig. 7). As we did not investigate the efferent connections of mPFC, it remains unclear which brain region is impacted by the reported disorganization of long-range inputs to deep layers of mPFC. However, existing literature indicated the involvement of mPFC projections to nucleus accumbens (NAc) in social approach behavior (63). Considering that the organization of long-range inputs in SS mice was impaired only in the deep layers of the mPFC, which predominantly project to subcortical regions, it is possible that imbalance in BLA and VH inputs to mPFC deep layers may disrupt the activity in mPFC-NAc circuit. This issue warrants further investigation in future studies.

**Figure 7:**
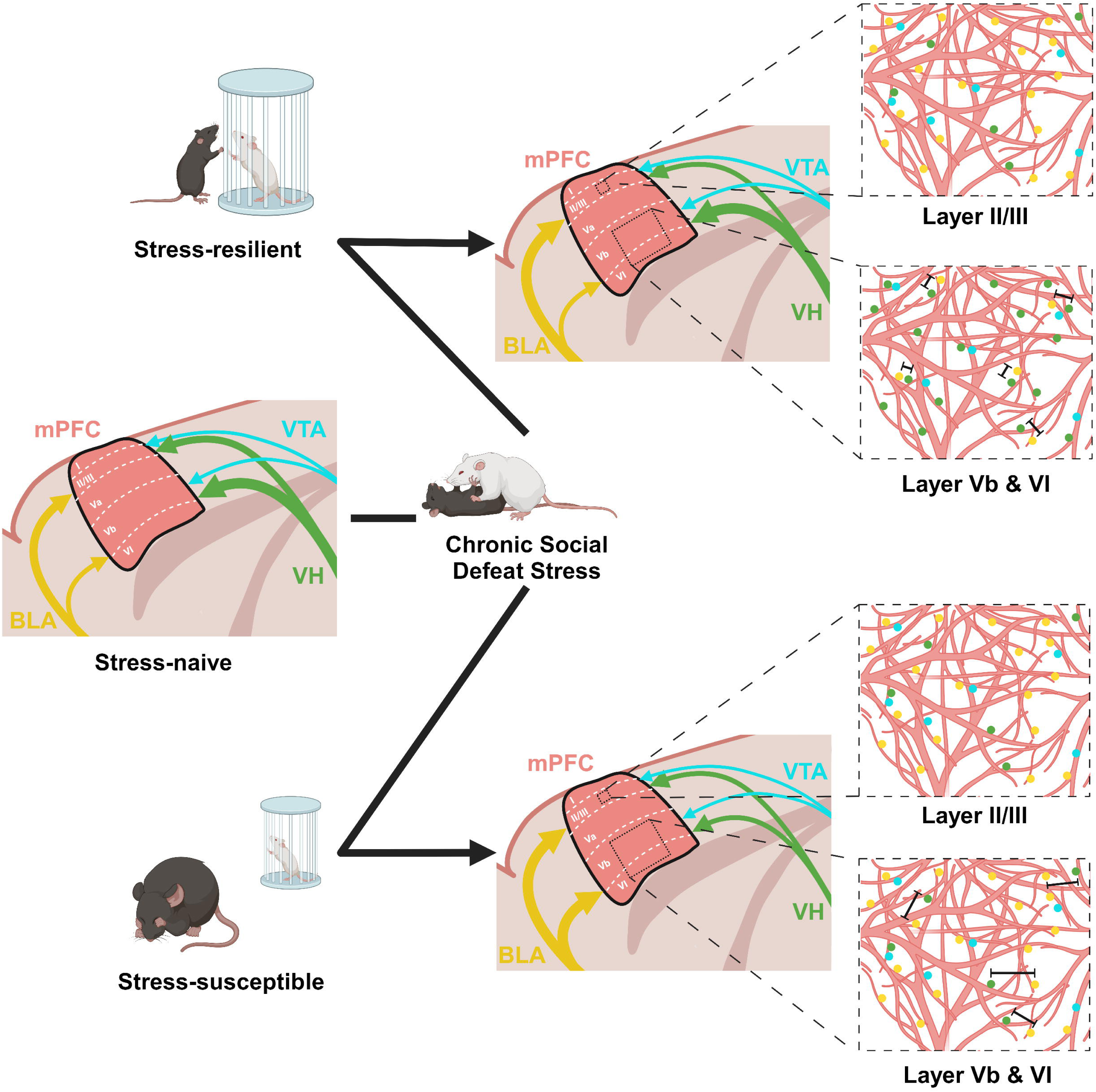
Summary scheme of the results. Chronic social defeat stress increases BLA-mPFC inputs in layer II/III, regardless of stress susceptibility. In contrast, in layers Vb&VI, it alters BLA-mPFC and VH-mPFC inputs only in SS mice. Note that the Euclidean distances between VH and their nearest BLA synapses were closer in SN&SR mice than that of SS mice (shown by black line segments).

How excitatory inputs from different brain regions onto mPFC contribute to different stress phenotypes is an intriguing question. One possibility is that these inputs may selectively target specific neuronal ensembles, each with distinct roles in fear-related behavior. Supporting this suggestion, our findings showed that inputs from the same brain region tend to make synapses on the same dendritic segments in SN and SR mice. In SS mice, however, we observed increase in the number of inputs from different brain regions converging onto the same dendritic segment. Additionally, our data indicates that under normal physiological conditions, inputs from the BLA typically terminate in close proximity to VH inputs. However, it seems essential for some VH-mPFC synapses to be positioned at a greater distance from BLA-mPFC synapses. In SS mice, this innervation pattern is disrupted. One may speculate that proximity of BLA inputs to VH inputs play a role in integrating threat-signals into ongoing information processing within those neuronal ensembles, while VH inputs distant from BLA inputs may be involved in processing non-threat-related information. Whether these distinct inputs originate from distinct mPFC-projecting VH neuronal populations, each with opposite roles on fear-related behaviors, remains to be addressed in future studies (25). Interestingly the disorganization in VH and BLA innervation of deep mPFC layers was not evident at the level of single dendritic segments, as there were no differences in the percentage of VH and BLA inputs on single dendritic segments among all three groups.

Most of the earlier studies on the role of VTA on stress response that focused on dopaminergic neurons have indicated the role of VTA in regulating both stress response and social behavior. Although the ratio of VTA-to all synaptic inputs [VTA/(BLA+VH+VTA)] was similar across all cortical layers and groups, we observed that in dendritic segments receiving input from multiple brain regions, changes in VTA input-containing synaptic pairs correlated with stress susceptibility. This suggests that VTA inputs also play a role in the regulation of stress response.

Our study primarily focused on long-range projections to pyramidal mPFC neurons, and we did not investigate projections terminating on mPFC interneurons. This decision was influenced by the challenges associated with confocal multicolor imaging, particularly the quantitative limit for visualizing multiple fluorophores simultaneously. Given various reports of altered excitatory:inhibitory balance in stress-related mental disorders long-range projections on mPFC interneurons may also contribute to fine-tuning the stress response (64–68). To provide a more comprehensive understanding of the neural circuitry underlying stress response and resilience, investigation of the role of long-range projections to mPFC interneurons is warranted in future studies.

Our study was conducted only in male mice, thereby restricting the generalizability of our findings to females. CSDS is generally used in identification of stress susceptibility in male mice due to challenges in implementing it in females. Resident mice of either sex do not attack the intruder females, so, unlike male mice, females do not naturally face such social attacks. Although a CSDS paradigm has been developed to induce social defeat in females by applying another CD1 male mouse’s urine to vaginal orifices of female intruders (69), we chose to conduct our study in male mice to explore the mechanisms underlying stress susceptibility and resilience, given its higher ethological validity in males.

The sample size is limited owing to the rigorous nature of the imaging. To address this limitation, we performed principal component analysis and random forest classification, which confirmed that the observed changes are unlikely to be due to chance.

The computational tools that enable the large-scale 3D synaptic study pursued here have some limitations. While automated tracing of dendrites broadly succeeds, it can introduce mistakes, especially by linking segments that belong to different neurons (32). We addressed it by using a higher threshold for segmentation. However, this resulted in shorter segments, affecting the calculation of dendritic path distances (e.g., to the closest synapses). Therefore, instead, we focused on the Euclidean distances between synapses. Not only is this calculation very robust, but it also provides information about the 3D spatial organization of VH-mPFC, VTA-mPFC, and BLA-mPFC synapses across a column of mPFC.

In conclusion, the findings of the current study enhance our comprehension of chronic stress effects on brain circuitry in both physiological and pathological states by focusing on the synapses formed by long-range projections on mPFC. We have demonstrated laminae- and afferent-specific changes in stressed groups depending on their stress phenotype (resilient vs susceptible), highlighting potential avenues for developing preventive or therapeutic strategies in stress susceptible individuals following a stressful or traumatic event.

## Key resources table

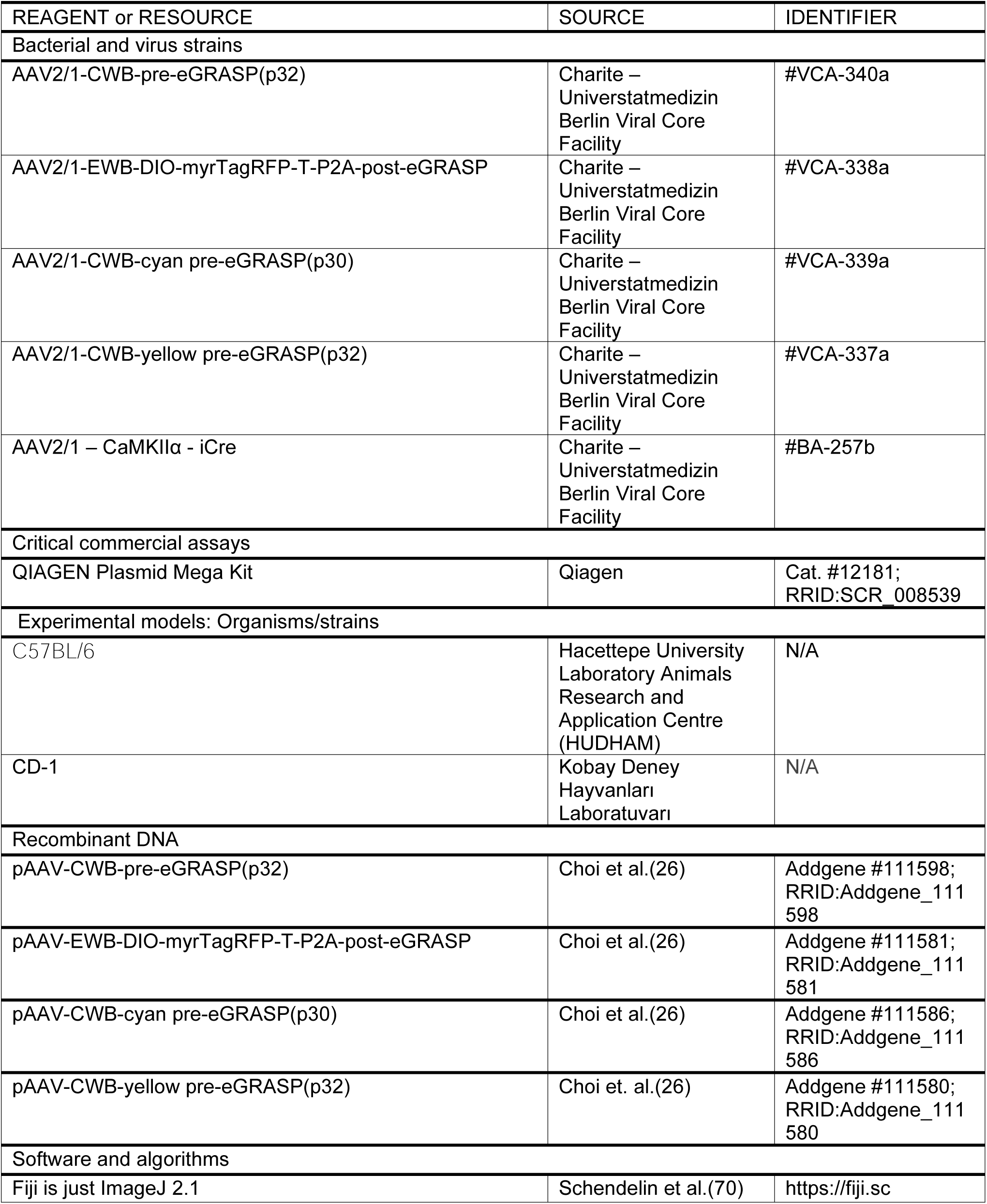
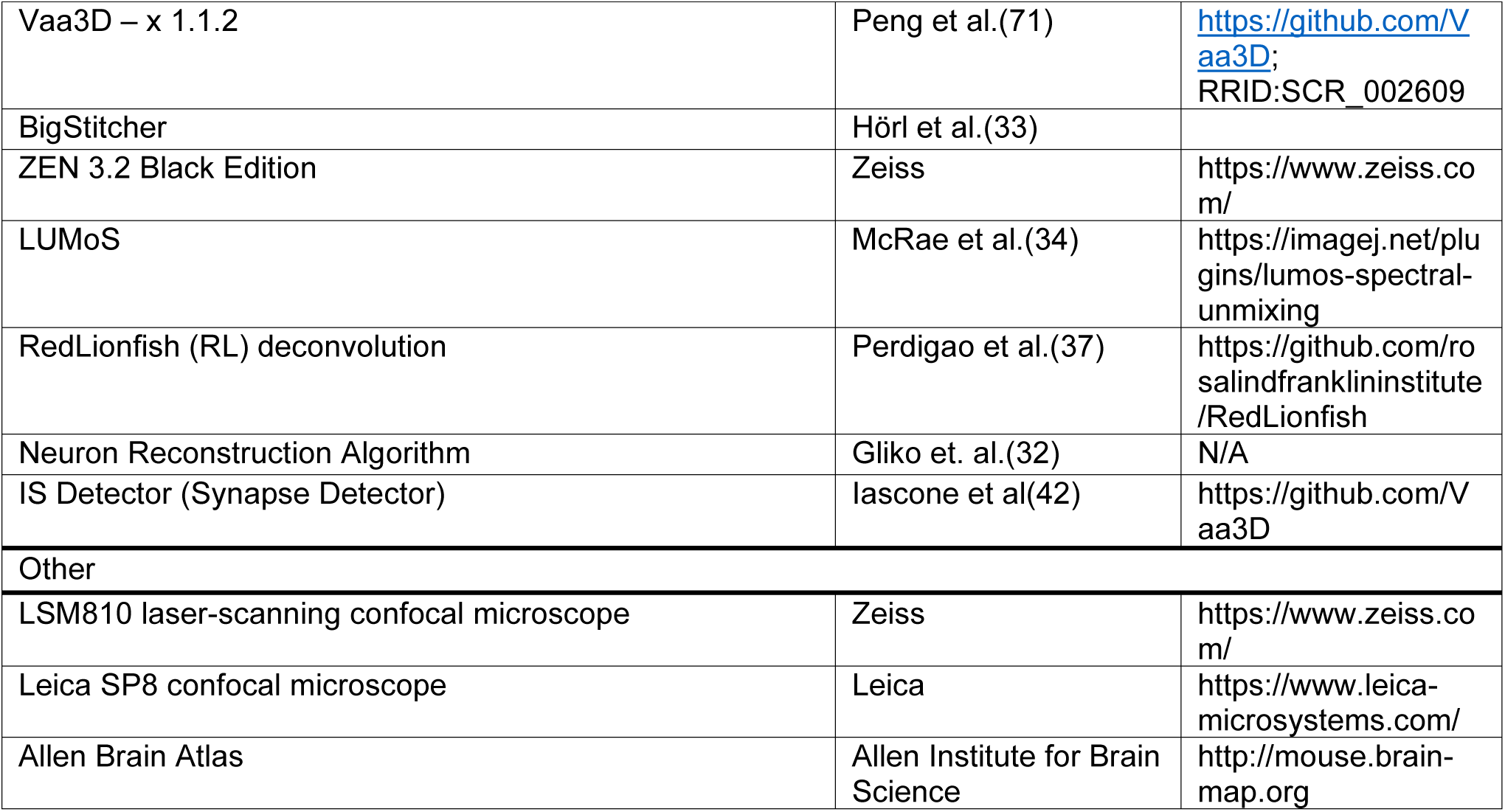

## AUTHOR CONTRIBUTIONS

STS, SEE and EEK were responsible for the design and conception of the study. STS, SEE and AC were responsible for the acquisition of data, STS, OG, SEE, AC, US and EEK were responsible for the analysis and interpretation of data. STS, US and EEK drafted the manuscript.

## ACKNOWLEDGEMENTS

This study is supported by Research Fund of Hacettepe University, Grant number: TSA-2020-18753 to EEK. We thank Dr. Turgay Dalkara for his valuable opinions on the manuscript; Dr. Nevin Belder, Dr. Aslıhan Bahadır-Varol, Mesut Fırat and Anke Schönherr for their technical support. Graphical depictions inside Figure-1, Figure-2, Figure-6 and Figure 7 were prepared with Biorender.

---

**Supplemental Table 1.**
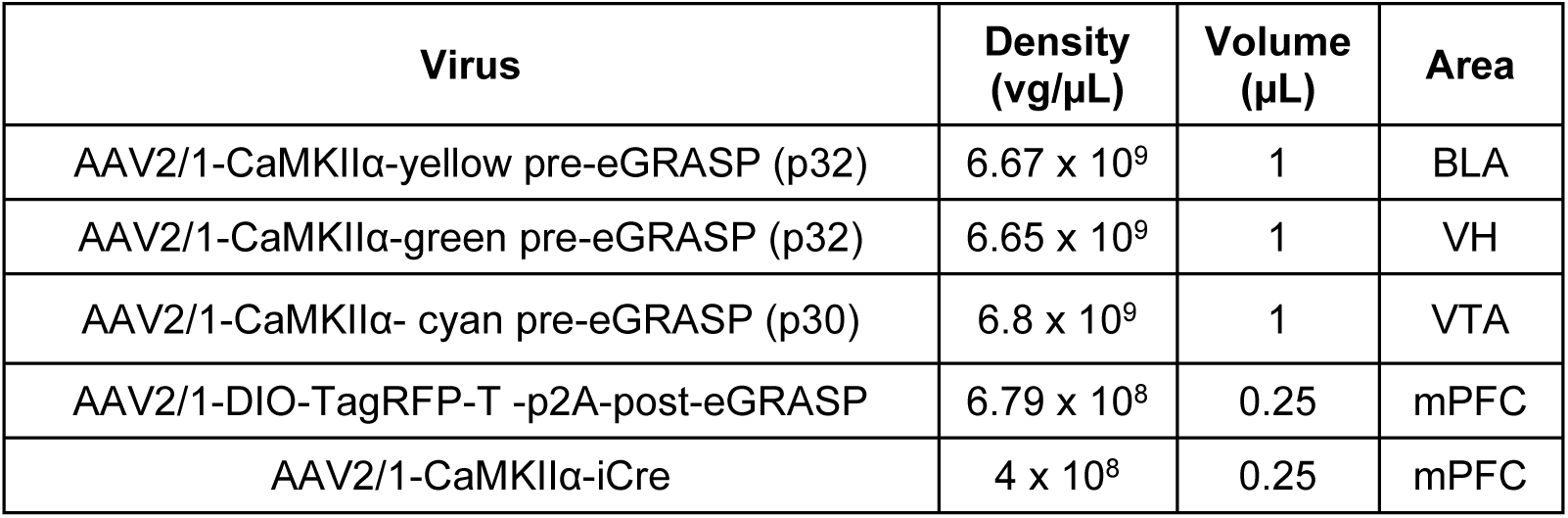
Characteristics, injection sites and volumes of the adeno-associated-virus.

| **Virus** | **Density**<br>**(vg/μL)** | **Volume**<br>**(μL)** | **Area** |
| --- | --- | --- | --- |
| AAV2/1-CaMKIIα-yellow pre-eGRASP (p32) | 6.67 x 10<sup>9</sup> | 1 | BLA |
| AAV2/1-CaMKIIα-green pre-eGRASP (p32) | 6.65 x 10<sup>9</sup> | 1 | VH |
| AAV2/1-CaMKIIα- cyan pre-eGRASP (p30) | 6.8 x 10<sup>9</sup> | 1 | VTA |
| AAV2/1-DIO-TagRFP-T -p2A-post-eGRASP | 6.79 x 10<sup>8</sup> | 0.25 | mPFC |
| AAV2/1-CaMKIIα-iCre | 4 x 10<sup>8</sup> | 0.25 | mPFC |

**Supplemental Table 2.** Results of Kruskal-Wallis and Mann-Whitney U test comparisons between Euclidean distances of synapses originating from the same brain region or from multiple brain regions in layer II/III and layer Vb & VI.

| Layers | Bins<br>(μm) | KW | MWU |  |  |  |
| --- | --- | --- | --- | --- | --- | --- |
|  |  |  | SN vs STR | SS vs SR | SS vs SN | SR vs SN |
| From VH to closest VH synapse (Fig. 6A and Suppl. Fig. 8A) |  |  |  |  |  |  |
| Layer<br>II/III | 0-5 | 0.67 | 1.00 |  | N/A |  |
|  | 5-10 | 0.98 | 0.95 |  | N/A |  |
|  | 10-15 | 0.24 | 0.32 |  | N/A |  |
|  | 15-20 | **0.056** | **0.022** | **0.51** | **0.059** | **0.059** |
|  | 20-25 | 0.55 | 0.67 |  | N/A |  |
|  | 25-30 | 0.73 | 0.84 |  | N/A |  |
|  | 30-35 | 0.55 | 0.31 |  | N/A |  |
|  | 35+ | 0.35 | 0.66 |  | N/A |  |
| Layer<br>Vb & VI | 0-5 | 0.29 | 1.00 |  | N/A |  |
|  | 5-10 | 0.45 | 0.25 |  | N/A |  |
|  | 10-15 | 0.53 | 0.76 |  | N/A |  |
|  | 15-20 | 0.26 | 0.95 |  | N/A |  |
|  | 20-25 | 0.29 | 0.80 |  | N/A |  |
|  | 25-30 | 0.80 | 0.95 |  | N/A |  |
|  | 30-35 | 0.24 | 0.62 |  | N/A |  |
|  | 35+ | 0.15 | 0.75 |  | N/A |  |
| From BLA to closest BLA synapse (Fig. 6B and Suppl. Fig. 8B) |  |  |  |  |  |  |
| Layer<br>II/III | 0-5 | 0.20 | 0.08 |  | N/A |  |
|  | 5-10 | 0.69 | 0.53 |  | N/A |  |
|  | 10-15 | 0.72 | 0.80 |  | N/A |  |
|  | 15-20 | 0.37 | 0.20 |  | N/A |  |
|  | 20-25 | 0.48 | 0.25 |  | N/A |  |
|  | 25-30 | 0.39 | 0.24 |  | N/A |  |
|  | 30-35 | 0.31 | 0.22 |  | N/A |  |
|  | 35+ | 0.86 | 0.64 |  | N/A |  |
| Layer<br>Vb & VI | 0-5 | 0.22 | 0.09 |  | N/A |  |
|  | 5-10 | 0.28 | 0.24 |  | N/A |  |
|  | 10-15 | 0.67 | 0.46 |  | N/A |  |
|  | 15-20 | 0.96 | 0.85 |  | N/A |  |
|  | 20-25 | 0.40 | 0.21 |  | N/A |  |
|  | 25-30 | 0.81 | 0.71 |  | N/A |  |
|  | 30-35 | 0.53 | 0.38 |  | N/A |  |
|  | 35+ | 0.18 | 0.12 |  | N/A |  |

| Layers | Bins<br>($\mu\text{m}$) | KW | MWU | | | |
| --- | --- | --- | --- | --- | --- | --- |
|  |  |  | SN vs STR | SS vs SR | SS vs SN | SR vs SN |
| From VH to closest BLA synapse (Fig. 6C and Suppl. Fig. 8C) |  |  |  |  |  |  |
| Layer II/III | 0-5 | 0.06 | 0.055 | 0.42 | 0.54 | 0.0079 |
|  | 5-10 | 0.94 | 0.80 |  | N/A |  |
|  | 10-15 | 0.27 | 0.16 |  | N/A |  |
|  | 15-20 | 0.12 | 0.13 |  | N/A |  |
|  | 20-25 | **0.022** | **0.0086** | **0.42** | **0.07** | **0.025** |
|  | 25-30 | **0.031** | **0.013** | **0.42** | **0.11** | **0.025** |
|  | 30-35 | **0.022** | **0.0086** | **0.42** | **0.07** | **0.025** |
|  | 35+ | 0.13 | 0.06 |  | N/A |  |
| Layer Vb & VI | 0-5 | **0.009** | **0.027** | **0.015** | **0.0079** | **0.30** |
|  | 5-10 | 0.35 | 0.30 |  | N/A |  |
|  | 10-15 | 0.10 | 1.00 |  | N/A |  |
|  | 15-20 | **0.034** | **0.37** | **0.031** | **0.031** | **0.69** |
|  | 20-25 | 0.14 | 0.50 |  | N/A |  |
|  | 25-30 | 0.06 | 0.07 | 0.13 | 0.034 | 0.42 |
|  | 30-35 | **0.036** | **0.055** | **0.16** | **0.011** | **0.54** |
|  | 35+ | 0.06 | 0.041 | 0.33 | 0.034 | 0.22 |
| From BLA to closest VH synapse (Fig. 6D and Suppl. Fig. 8D) |  |  |  |  |  |  |
| Layer II/III | 0-5 | 0.19 | 0.11 |  | N/A |  |
|  | 5-10 | 0.92 | 0.75 |  | N/A |  |
|  | 10-15 | 0.43 | 0.21 |  | N/A |  |
|  | 15-20 | 0.24 | 0.25 |  | N/A |  |
|  | 20-25 | 0.88 | 0.71 |  | N/A |  |
|  | 25-30 | 0.73 | 0.49 |  | N/A |  |
|  | 30-35 | 0.41 | 0.94 |  | N/A |  |
|  | 35+ | 0.53 | 0.53 |  | N/A |  |
| Layer Vb & VI | 0-5 | **0.004** | **0.00066** | **0.055** | **0.0079** | **0.0079** |
|  | 5-10 | **0.045** | **0.012** | **0.84** | **0.09** | **0.015** |
|  | 10-15 | **0.016** | **0.031** | **0.031** | **0.021** | **0.20** |
|  | 15-20 | 0.07 | 0.06 | 0.30 | 0.034 | 0.33 |
|  | 20-25 | **0.004** | **0.045** | **0.017** | **0.0074** | **0.42** |
|  | 25-30 | **0.046** | **0.045** | **0.23** | **0.025** | **0.17** |
|  | 30-35 | **0.031** | **0.08** | **0.11** | **0.025** | **0.42** |
|  | 35+ | 0.10 | 0.13 |  | N/A |  |

