## Supplementary figures and images for "Layer-specific input to medial prefrontal cortex is linked to stress susceptibility"

### Supplemental Figure - 1

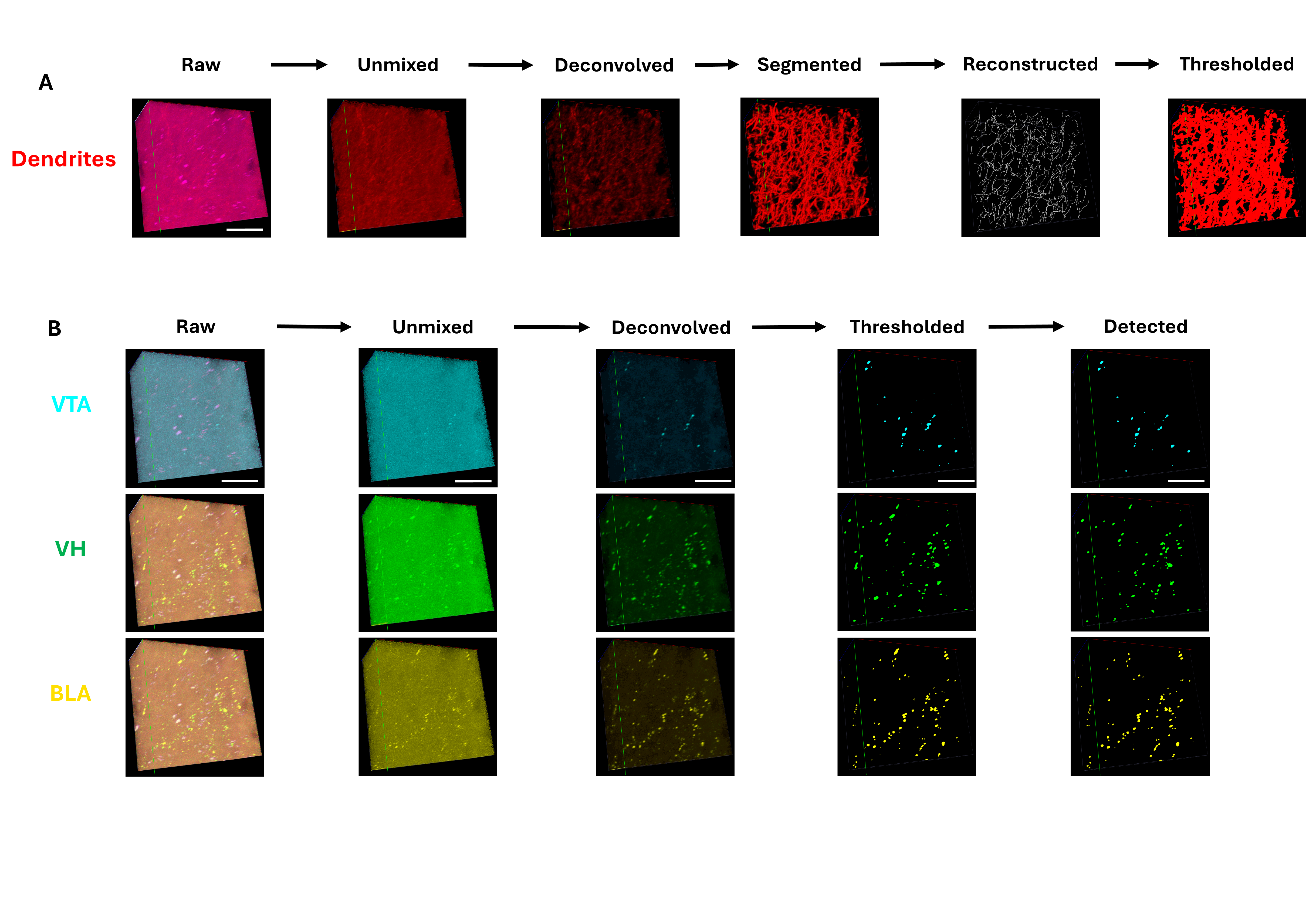

### Supplemental Figure - 2

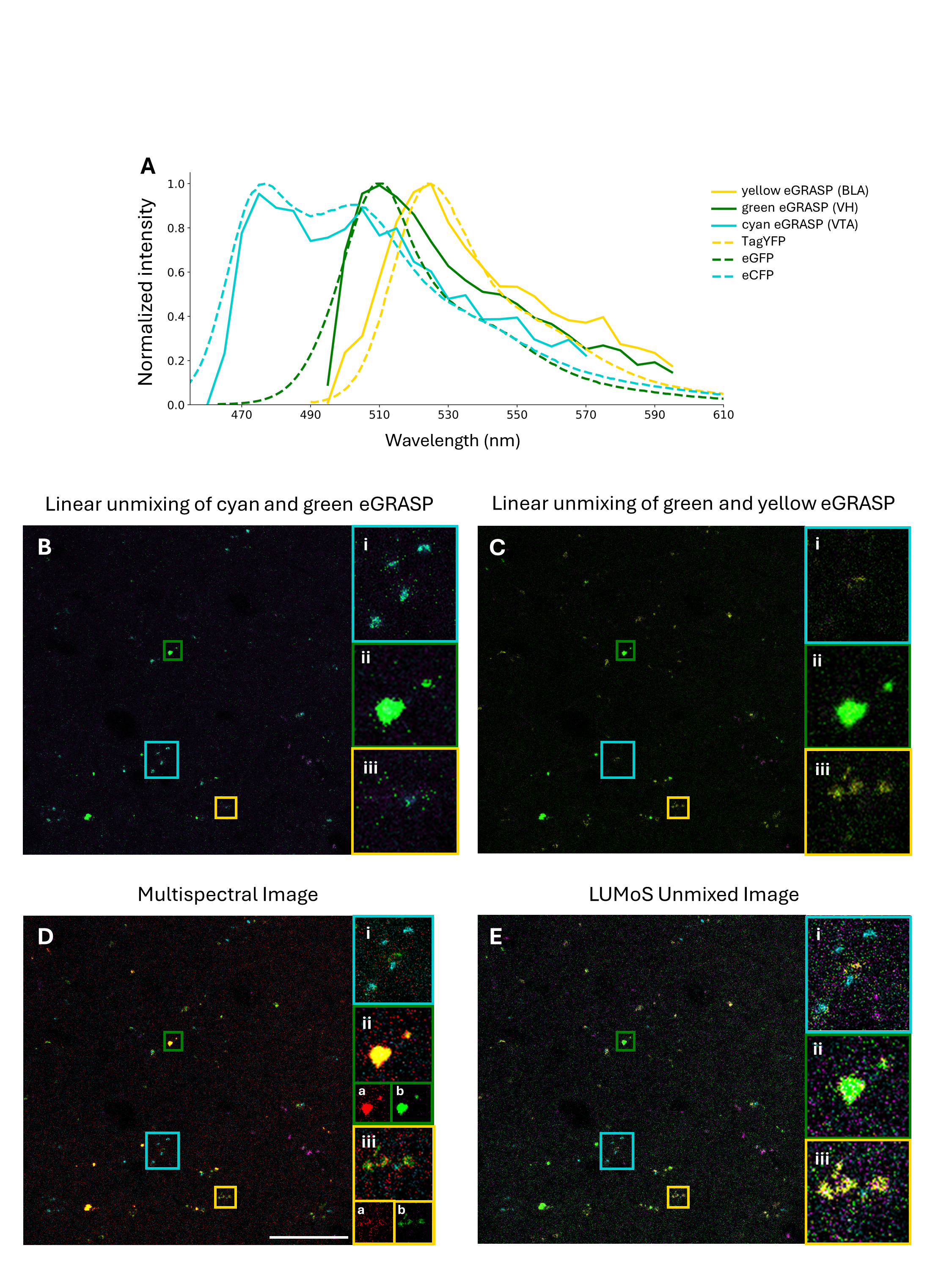

### Supplemental Figure - 3

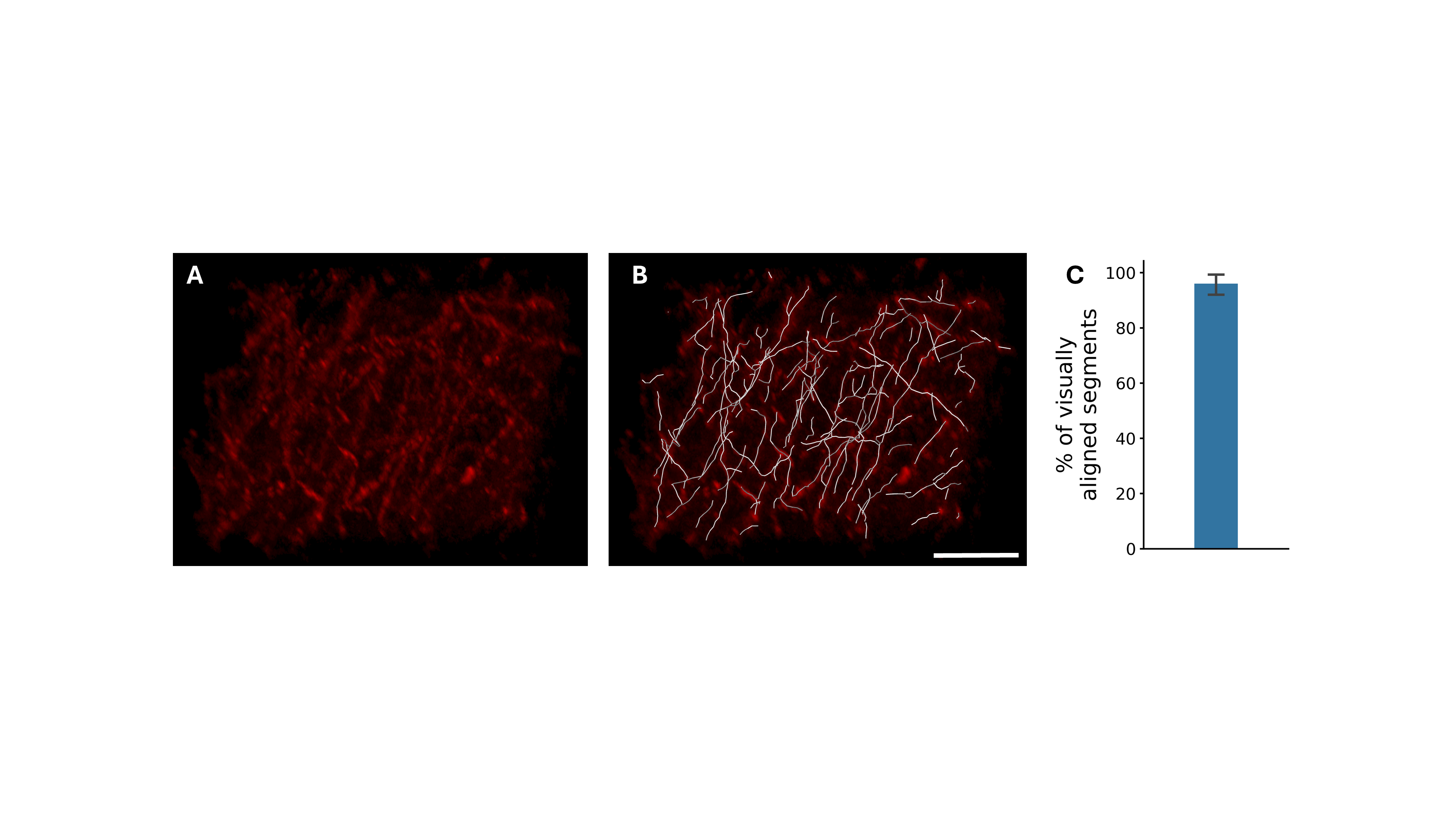

### Supplemental Figure - 4

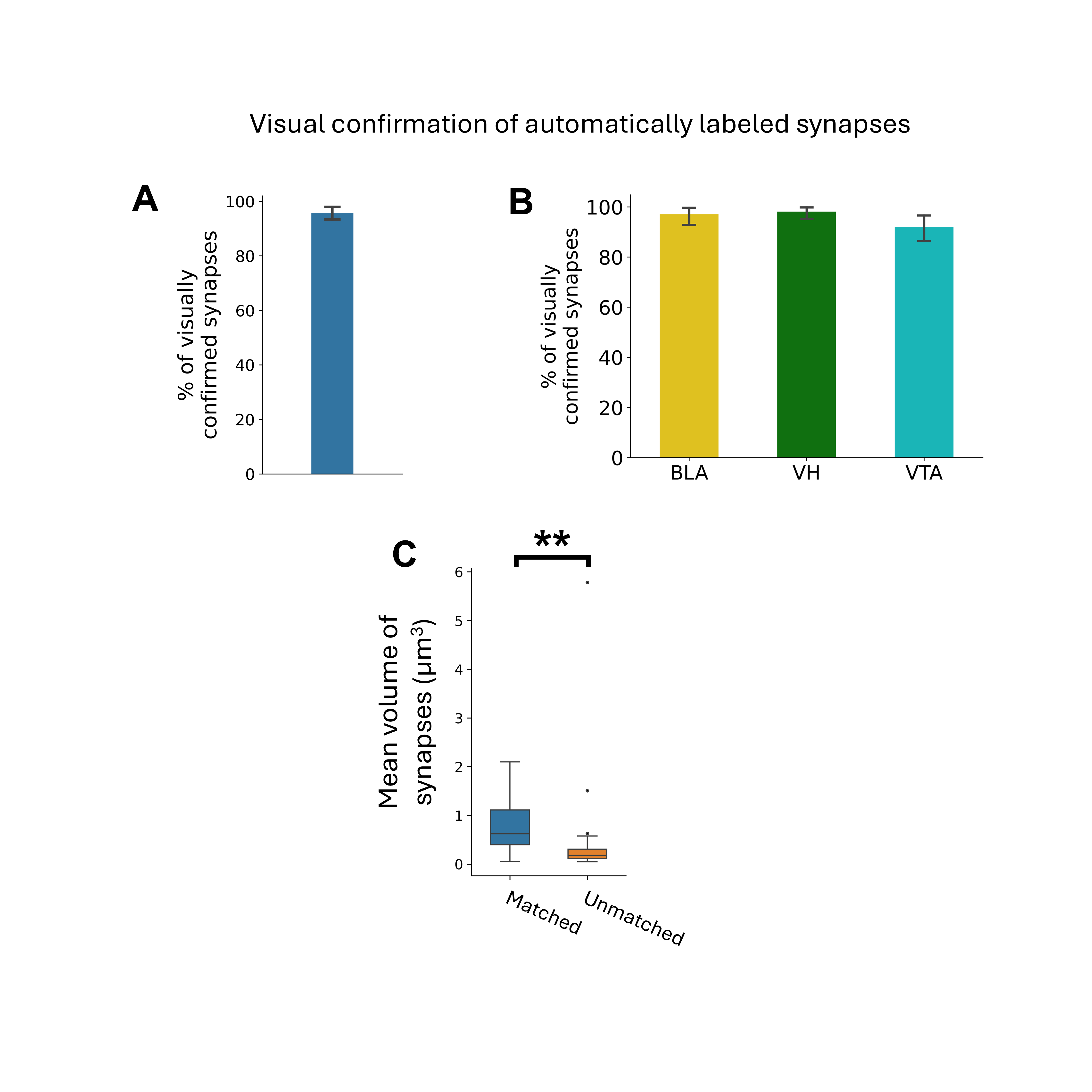

### Supplemental Figure - 5

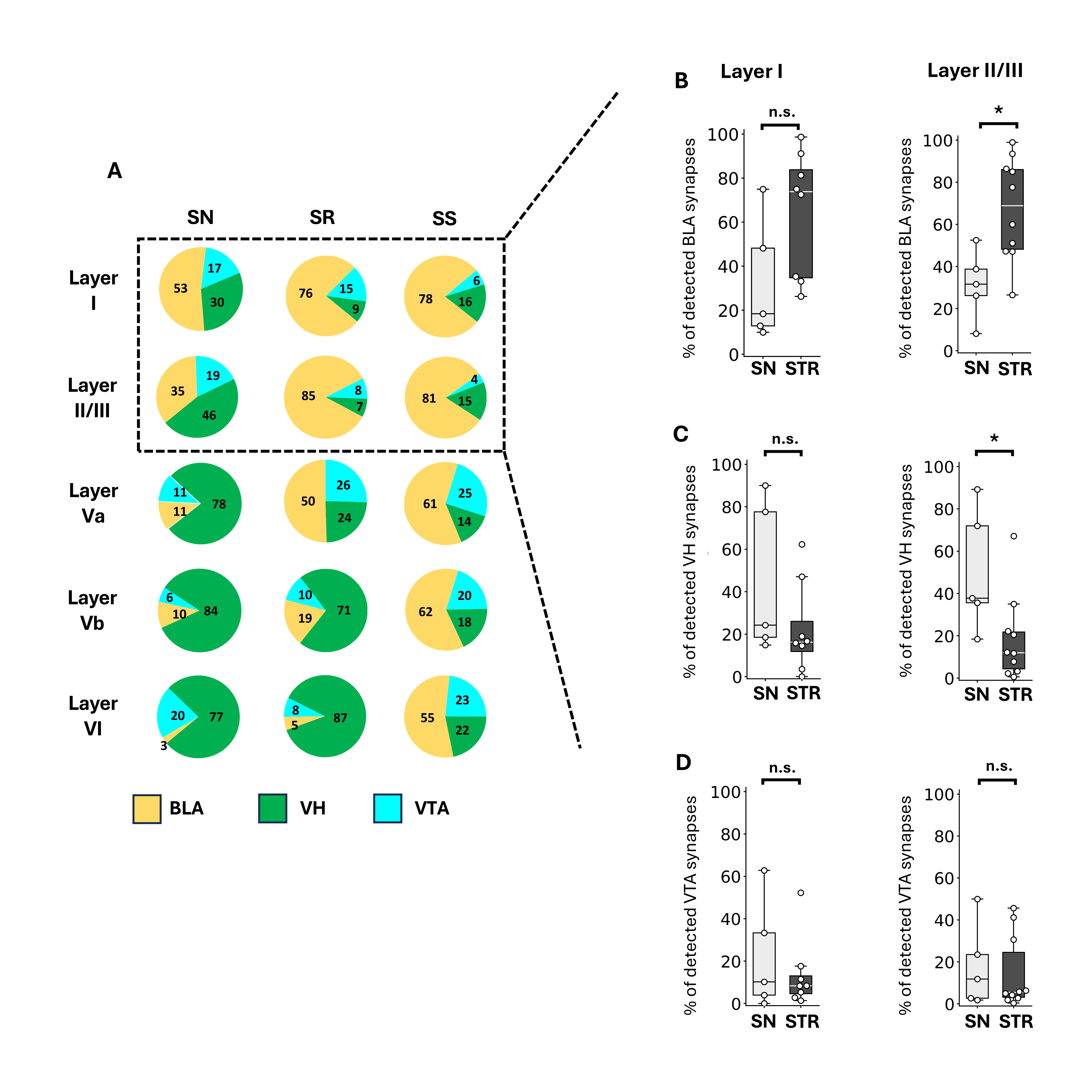

### Supplemental Figure - 6

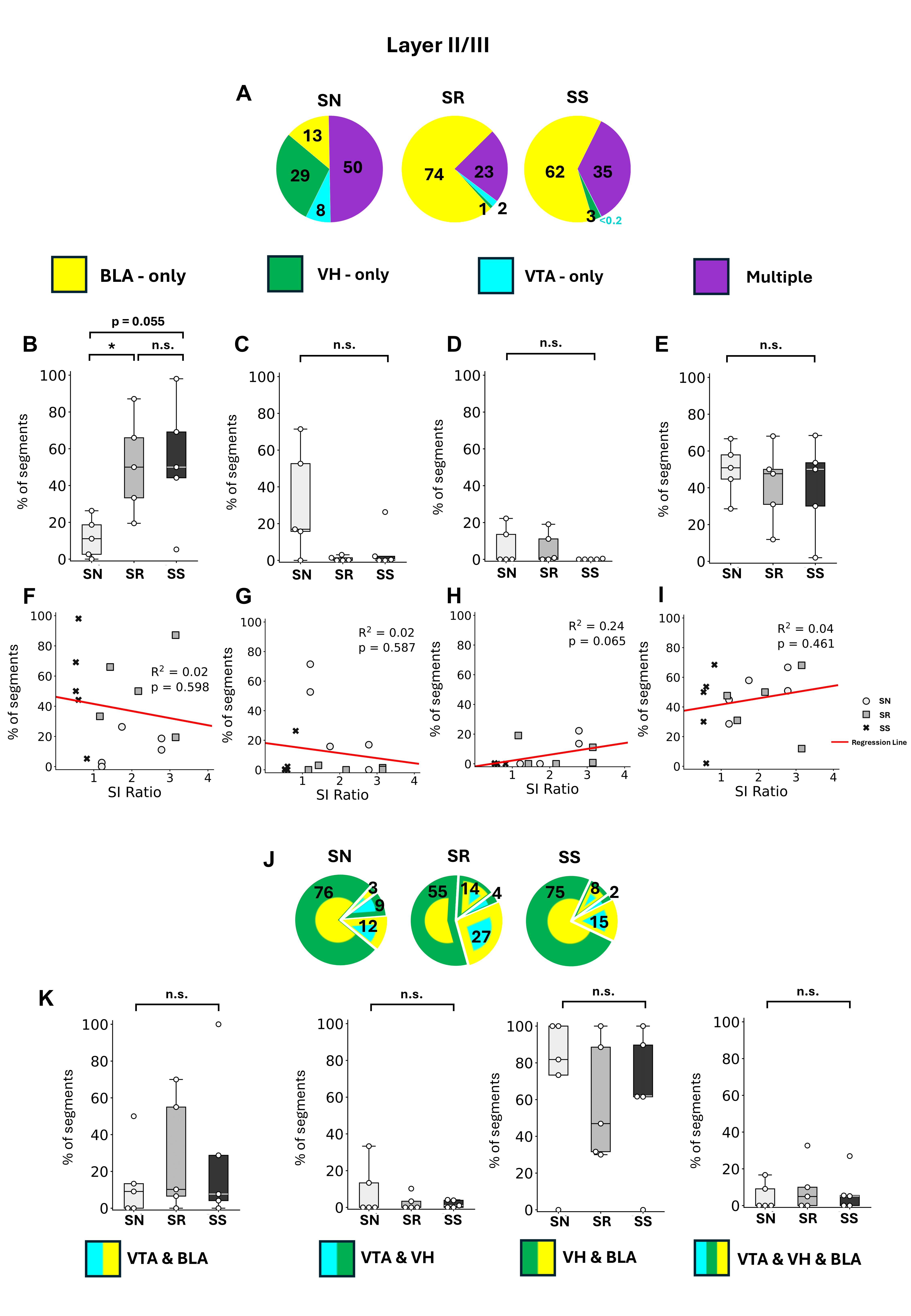

### Supplemental Figure - 7

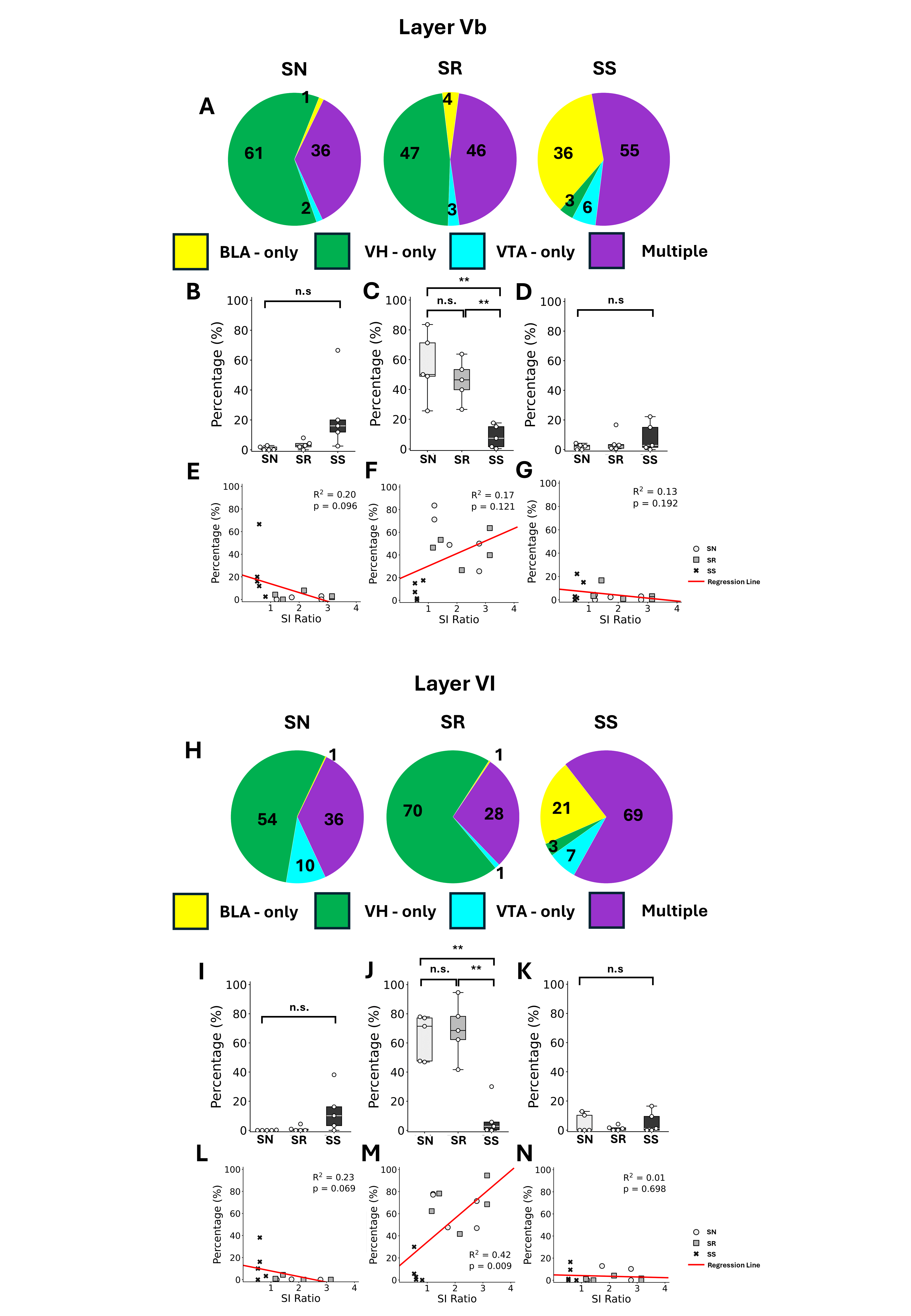

### Supplemental Figure - 8

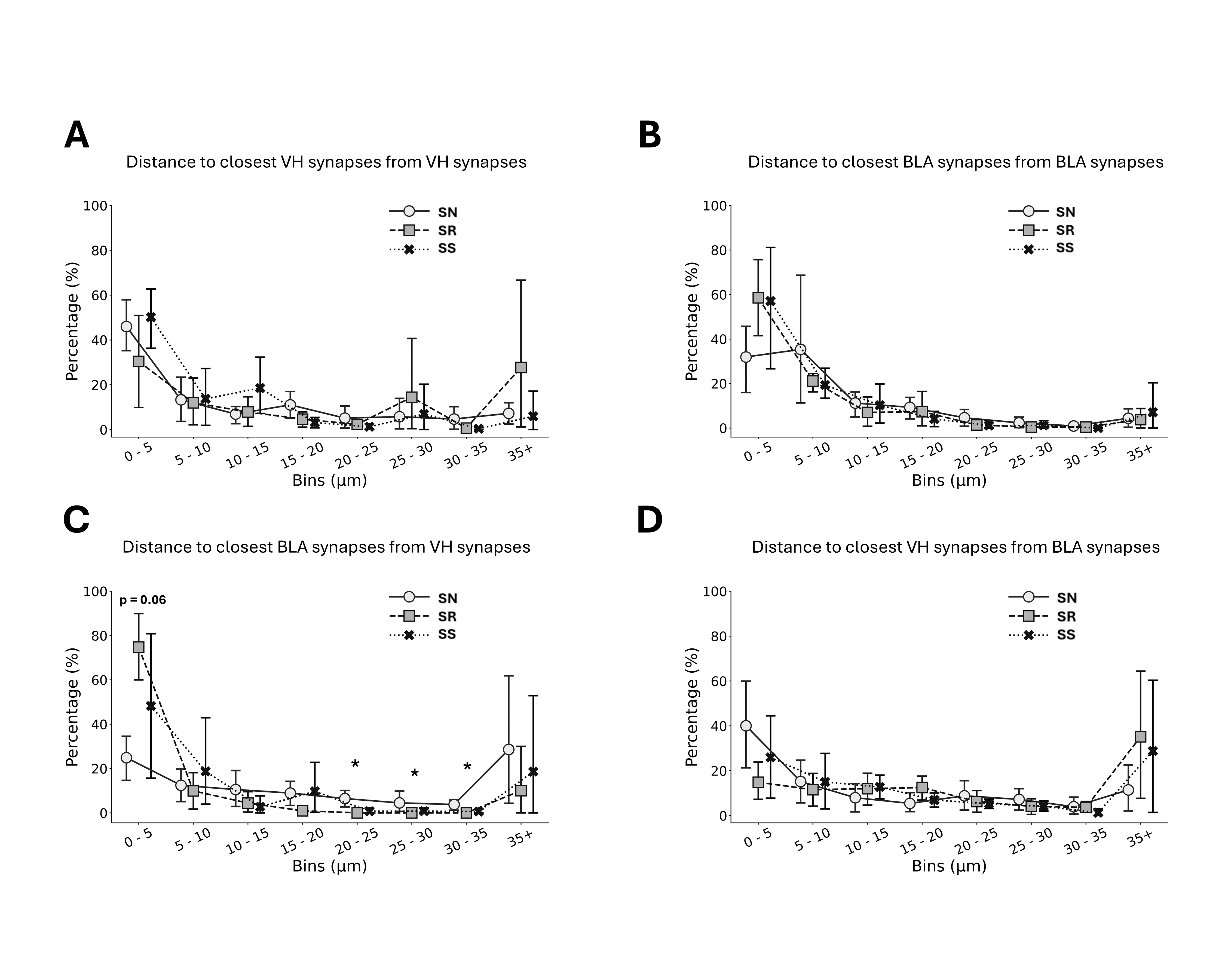
